# GIP receptor reduces osteoclast activity and improves osteoblast survival by activating multiple signaling pathways

**DOI:** 10.1101/2022.07.02.498420

**Authors:** Morten S. Hansen, Kent Søe, Line L. Christensen, Paula Fernandez-Guerra, Nina W. Hansen, Rachael A. Wyatt, Claire Martin, Rowan S. Hardy, Thomas L. Andersen, Jacob B. Olesen, Søren Overgaard, Bolette Hartmann, Mette M. Rosenkilde, Moustapha Kassem, Alexander Rauch, Caroline M. Gorvin, Morten Frost

**Author notes:** Correspondence should be addressed to: Caroline Gorvin or Morten Frost. These authors contributed equally to this manuscript and should be considered joint senior authors (Caroline M. Gorvin, Morten Frost).

## Abstract

Bone is a dynamic tissue that is remodeled throughout life by bone resorbing osteoclasts and bone forming osteoblasts, to adapt to physiological or mechanical demands. These processes are impaired in osteoporosis, and understanding how bone remodeling is regulated could improve anti-osteoporotic treatments. Clinical investigations show that short-term treatment with glucose-dependent insulinotropic polypeptide (GIP) acutely decreases serum markers of bone resorption and may increase bone formation. However, evidence for direct effects of GIP intracellular signaling and functions in mature human osteoclasts and osteoblasts have not been investigated. We report that the GIP receptor (GIPR) is robustly expressed in mature human osteoclasts. Exposure of osteoclasts to GIP inhibits osteoclastogenesis, delays bone resorption, and increases osteoclast apoptosis by acting upon multiple signaling pathways (cAMP, Src, Akt, calcium, p38) to impair nuclear translocation of nuclear factor of activated T cells 1 (NFATc1) and nuclear factor-κB (NFκB). Human osteoblasts also express GIPR, and GIP improves osteoblast survival via cAMP and Akt-mediated pathways. GIP treatment of co-cultures of osteoclasts and osteoblasts also decreased bone resorption. Antagonizing GIPR with GIP(3-30)NH_2_ abolished the effects of GIP on osteoclasts and osteoblasts. This study demonstrates that GIP inhibits bone resorption and improves survival of human osteoblasts, which could increase bone mass and strength, supporting clinical investigations of the effect of GIP on bone. Moreover, this study demonstrates that GIPR agonism could be beneficial in the treatment of disorders of bone remodeling, such as osteoporosis.

**One-sentence Summary:** GIP acts directly on bone cells to regulate bone remodeling

## Introduction

Bone is a dynamic tissue that is remodeled throughout life by bone resorbing osteoclasts and bone forming osteoblasts, to adapt to physiological or mechanical demands. Bone remodeling aims to maintain bone homeostasis and is regulated by multiple hormones, including parathyroid hormone and growth hormone (*1*). Osteoporosis is caused by an imbalance in bone remodeling and is characterized by decreased bone mass, resulting in increased fracture risk and affects >10 million individuals in the United States and >27 million in Europe (*1, 2*). Deciphering the regulation of bone remodeling has paved the way for development of the currently available anti-osteoporotic treatments, which increase bone mass and reduce fracture risk by decreasing bone resorption, stimulating bone formation, or through dual effects on bone resorption and formation (*1*).

A growing body of evidence from rodent and human studies indicates that the gut-secreted hormone GIP (glucose-dependent insulinotropic polypeptide or gastric inhibitory polypeptide) regulates bone remodeling (*3*). Compared to wild-type mice, homozygous transgenic mice with a *Gip* truncation had reduced bone volume and an increased osteoclast surface (*4*). Reciprocally, *Gip* overexpression in mice increased bone mineral density (BMD), raised serum osteocalcin, a marker of osteoblast activity, and reduced osteoclast numbers and bone resorption markers (*5*).

GIP acts on the GIP receptor (GIPR), a G protein-coupled receptor (GPCR), to activate G_s_-mediated increases in cAMP and mediate its primary function at pancreatic β-cells, where it increases glucose-dependent insulin secretion, prior to GIP’s degradation by the dipeptidyl peptidase IV (DPP4) enzyme (*6*). Several studies have also investigated whether GIPR is expressed on osteoclasts and osteoblast-like cells, but findings are inconsistent. Weak GIPR mRNA and/or protein expression levels have been reported in human and rodent osteoblast-like cell-lines (hMSC-TERT, SaOS2, MG63, ROS17/2.8, MC3T3-E1), and exposure of these cells to GIP has been described to increase cAMP, intracellular calcium (Ca^2+^_i_) and p38 (*7–9*). In addition, GIPR mRNA expression has been detected in cultured primary human osteoclasts (*9*). However, signaling studies in rodent cell-lines (Raw264.7) showed that a stable GIP analogue reduced Ca^2+^_i_ signaling and bone resorption without activating cAMP (*10, 11*).

Reports on the bone phenotype of global *Gipr* knockout mice (*Gipr^-/-^*) are inconsistent, possibly due to variations in the transgenic strategies used to generate the two models. *Gipr^-/-^*mice with deletion of *GIPR* exons 1-6 have been described to have reduced BMD, decreased circulating markers of bone formation, and elevated osteoclast numbers and resorption markers (*12–14*); however, another murine model with deletion of *GIPR* exons 4-5 showed fewer osteoclasts, increased trabecular bone volume and more active osteoblasts, despite reductions in bone strength (*15*).

In humans, GIP infusion suppressed the bone resorption marker C-terminal telopeptide of type I collagen (CTX) in healthy (*16*), overweight (*17*), type 1 diabetic (*18*), and hypoparathyroid individuals (*9*). Moreover, GIP transiently increased the bone formation marker procollagen type 1 N propeptide (P1NP) in healthy men and type 1 diabetics (*18, 19*), but not in overweight individuals or patients with hypoparathyroidism (*9, 17*). Importantly, pre-treatment with GIP(3-30)NH_2_, a high-affinity GIPR antagonist (*20*), abolished GIP-induced CTX and P1NP responses in healthy men (*19*). Finally, a rare missense *GIPR* variant in humans (rs143430880) is associated with lower BMD, although overall bone fracture risk is not affected (*21*).

While murine models demonstrate that GIP and GIPR influence bone mass and strength, and clinical studies support that GIP acutely regulates bone resorption and bone formation, direct effects of GIP treatment and GIPR antagonism on intracellular signaling and functions in mature human osteoclasts and osteoblasts have not been investigated. Therefore, we conducted a comprehensive analysis of the effect of GIP on GIPR-mediated signaling in primary human osteoclasts and osteoblasts. Robust expression of *GIPR* was demonstrated in human osteoclasts. By utilization of highly sensitive quantitative assays combined with specific antagonists of the GIPR and signal components, we showed that GIP acts upon multiple signaling pathways to reduce nuclear translocation of nuclear factor of activated T cells 1 (NFATc1) and nuclear factor-κB (NFκB), impairs transcription of genes involved in osteoclast survival and differentiation, increases apoptosis, and decreases bone resorption. Moreover, we confirmed *GIPR* expression in human osteoblasts and showed that GIP activates cAMP and Akt pathways and improves cell survival. Finally, GIP inhibited bone resorption without affecting mineralization in osteoblast and osteoclast co-cultures.

## Results

### GIP reduces bone resorption by direct action on osteoclast GIPR

Previous studies have shown that *GIPR* is expressed on human osteoclasts (*9*) and that exposure to a GIP analogue during osteoclastogenesis reduces resorptive activity of mature osteoclasts (*11*). We confirmed the expression of GIPR on mature osteoclasts by *in situ* hybridization (Figure 1A) and showed that *GIPR* mRNA expression increases during osteoclastogenesis (>13-fold increase from day 1 to day 10 of differentiation) (Figure 1B) on cells differentiated from human CD14^+^ monocytes.

**Figure 1.**
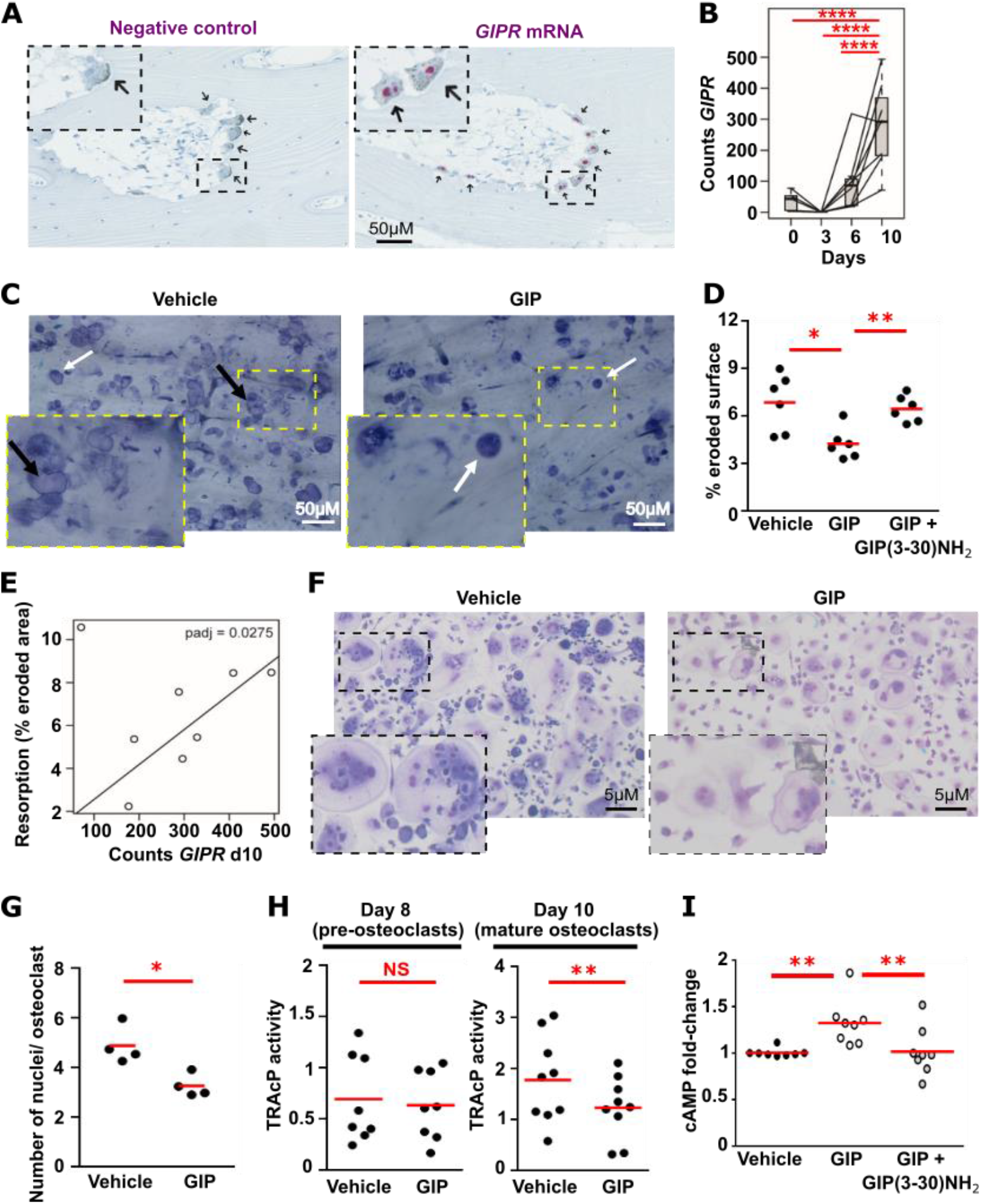
GIP reduces bone resorption and activates cAMP signaling in primary human osteoclasts. (**A**) *In situ* hybridization showing *GIPR* mRNA expression (magenta) in human osteoclasts (arrows). (**B**) Quantification of *GIPR* mRNA expression during osteoclast differentiation by RNA-sequencing. N=8 donors. DEseq2 adjusted p-values modeled for donor and timepoint. (**C**) Representative images and (**D**) quantification of resorption pits and trenches formed on bone slices by osteoclasts exposed to vehicle or 10nM GIP, visualized using toluidine blue. N=6 donors. White arrows show pits and black arrows show trenches. (**E**) Baseline resorptive activity of osteoclasts over *GIPR* mRNA expression at day 10 of osteoclast differentiation. N=8 donors. DEseq2 adjusted p-values modeled for resorptive activity. (**F**) Representative images and (**G**) quantification of number of nuclei per osteoclast visualized using May-Grünwald-Giemsa stain. N=4 donors. (**H**) Quantification of osteoclast activity measured by TRAcP staining. Osteoclasts were exposed to vehicle, 10nM GIP or GIP with the GIPR antagonist, GIP(3-30)NH_2_, for 3 days in panels C-H. (**I**) Quantification of cAMP generated in osteoclasts exposed to vehicle, GIP or GIP+GIP(3-30)NH_2_ for 30 minutes measured by LANCE assay. Each dot represents one donor with mean or median shown in red in panels D, G, H, I. ****p<0.0001, **p<0.01, *p<0.05. Statistical analyses were performed by: one-way ANOVA with Dunnett’s correction for multiple comparison testing for panel 1D; paired t-test for panels 1G and 1H; and Kruskal-Wallis one-way ANOVA with Dunn’s multiple comparisons test for 1I.

To confirm that GIP affects bone resorption we performed 3 analyses: measurement of osteoclastic resorptive activity by assessing percentage eroded surface on bovine bone slices, quantification of the number of nuclei in mature osteoclasts, and assessment of tartrate-resistant acid phosphatase 5b (TRAcP) activity. First, GIP decreased bone resorption in osteoclast monocultures exposed to increasing concentrations of GIP (0.1, 1, 10, 100 nM) for 3 days, when compared to vehicle-treated cells (Figure 1C and S1A). The effects of GIP on bone resorption were also observed in co-cultures of human osteoclasts with osteoblastic (immature-like reversal) cells (Figure S1B), demonstrating that GIP-mediated reductions in bone resorption can occur in a more physiologically relevant system. Subsequent studies were performed using 10nM GIP. Pre-treatment of mature osteoclasts with a specific GIPR antagonist, 10µM GIP(3-30)NH_2_, prevented the GIP-mediated reductions in bone resorption (Figure 1D). Osteoclasts resorb bone by forming pits (round cavitas) or trenches (prolonged excavations) depending on the rate between collagenolysis versus demineralization (*22*). The ratio between pits and trenches was not affected by GIP, showing that GIP does not affect the bone resorption pattern. GIPR expression also correlated with the resorptive activity of mature osteoclasts (Figure 1E) Exposure to GIP during osteoclastogenesis reduced the number of nuclei per osteoclast (Figure 1F-G), and reduced TRAcP activity in human mature osteoclasts, but not in pre-osteoclasts (Figure 1H), consistent with induction of GIPR mRNA levels late during osteoclastogenesis. Taken together GIP has a profound inhibitory effect on late osteoclast differentiation and bone resorptive activity.

### GIPR activation enhances cAMP signaling in human osteoclasts

GIPR couples to Gs signaling pathways, which activate adenylate cyclase and increase cAMP. However, previous studies using mouse osteoclast-like cell-lines showed that a stable GIP analogue does not change cAMP levels (*11*). To determine whether GIP activates cAMP signaling pathways in human osteoclasts we measured cAMP accumulation by LANCE assays. We initially quantified cAMP accumulation in osteoclasts following exposure to either vehicle or GIP at several time points. GIP, but not vehicle, increased cAMP levels in osteoclasts after 30 minutes (Figure S1C). As exposure to GIP for 15 minutes, or more than 30 minutes, had no impact on cAMP levels, subsequent GIP signaling studies in osteoclasts were examined after 30 minutes GIP treatment. To confirm that GIP stimulation of osteoclasts was due to direct activation of GIPR, cells were pre-treated with GIP(3-30)NH_2_ for 60 minutes. GIP(3-30)NH_2_ pretreatment abolished GIP induced cAMP accumulation (Figure 1I). Increased cAMP was also observed in osteoclast-osteoblast co-cultures following 30 minutes exposure to GIP, but not in those pre-treated with GIP(3-30)NH_2_ (Figure S1D).

Stimulation of cAMP activates protein kinase A (PKA) and can act on several downstream signaling pathways including: enhancement of the cAMP response element binding protein (CREB) pathway, which increases bone resorption (*23*); and inhibition of the NFATc1 pathway, which reduces bone resorption (*24*). To determine whether GIP activates CREB signaling, the accumulation of phosphorylated CREB (p-CREB) was assessed by AlphaLISA assays. This showed no significant difference between p-CREB concentrations in osteoclasts exposed to vehicle or GIP in monocultures or osteoclast-osteoblast co-cultures (Figure S1E-F). Thus, GIPR and cAMP likely modify other signaling pathways than phosphorylation of CREB in human osteoclasts.

### p-Src is decreased by GIPR activation in osteoclast monocultures

As GIP reduces bone resorption, we next assessed signaling pathways known to affect osteoclast activity. Activation of the tyrosine kinase c-Src by phosphorylation plays a central role in actin ring formation, a requirement for bone resorption (*25*), and disruption of *Src* in mice induces osteopetrosis, a disorder characterized by decreased bone resorption (*26*). To assess Src signaling, AlphaLISA assays that measure phosphorylation of the Src-Tyr419 residue were performed. Osteoclasts exposed to GIP had significantly less phosphorylated Src (p-Src) than vehicle-treated cells (Figure 2A). When pre-treated with GIP(3-30)NH_2_, no difference was observed between vehicle or GIP treated cells, indicating the reduction in p-Src is GIPR mediated. Similar effects of GIP on p-Src were observed in osteoclast-osteoblast co-cultures (Figure S2A).

**Figure 2.**
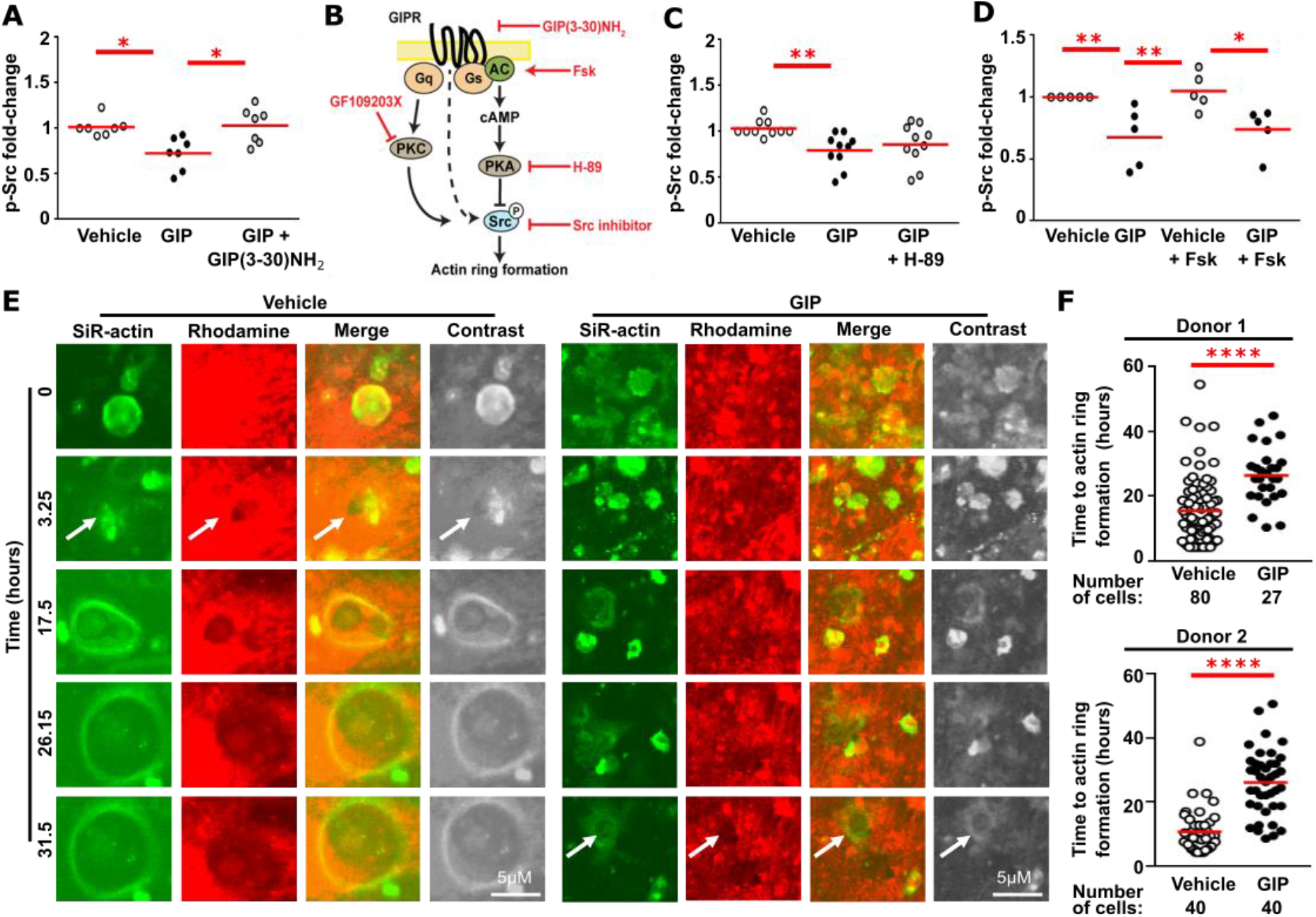
GIP reduces c-Src signaling and actin ring formation in human osteoclasts. (**A**) Quantification of phosphorylated c-Src (p-Src) in osteoclasts exposed to vehicle, GIP, or GIP with GIP(3-30)NH_2_ for 30 minutes measured by AlphaLISA. (**B**) Cartoon showing signaling pathways that may affect p-Src signaling. Inhibitors/ activators are in red. (**C-D**) Effect of pre-treatment with (C) the PKA inhibitor H-89 or (D) the adenylate cyclase activator forskolin (Fsk) on GIP-mediated p-Src responses in osteoclasts. p-Src was normalized to GAPDH in A, C, D. N=7 in A, N=10 in C, N=5 donors in D. Each point in A, C, D represents one donor, with mean or median shown in red. (**E**) Representative time-lapse images and (**F**) quantification of initiation of actin ring formation in 2 donors. Arrows indicate the actin ring. Each dot represents one cell measured with median shown in red. ****p<0.0001, **p<0.01, *p<0.05. Statistical analyses were performed by: Kruskal-Wallis one-way ANOVA with Dunn’s multiple comparisons test for 2A; one-way ANOVA with Sidak’s and Dunnett’s multiple comparisons tests, respectively, for panels 2B and 2C; nested t-test for 2F.

Elevations in cAMP can reduce Src activity in a PKA-dependent pathway (*27*). We therefore hypothesized that activation of GIPR-cAMP pathways could enhance PKA and reduce p-Src generation (Figure 2B). To test this, AlphaLISA assays were performed with the PKA inhibitor H-89. Pre-treatment of osteoclasts with H-89 for 60 minutes, followed by 30 minutes exposure to GIP or vehicle had no effect on p-Src (Figure 2C). To verify that disrupting the cAMP-PKA pathway does not affect GIP-mediated suppression of p-Src we repeated the assays with forskolin, which activates adenylate cyclase and should thus phenocopy any treatments that activate cAMP. Treatment with forskolin alone did not affect p-Src concentrations in osteoclasts, while a combined treatment with forskolin generated similar concentrations of p-Src to cells treated with vehicle (Figure 2D). Therefore, it is unlikely that the canonical GIPR-Gs-cAMP-PKA pathway modulates p-Src levels. Moreover, pre-treatment of osteoclasts with a PKC inhibitor (GF109203X) or a calcium chelator (BAPTA) had no effect on p-Src concentrations (Figure S2B). This indicates it is unlikely that GIPR couples to G_q_ pathways to activate p-Src. Additionally, inhibition of other signaling pathways including phosphatidylinositol-3-kinase (PI3K)-Akt, which are known to be activated by GIPR in other cells (*28*), had no effect on p-Src, although a Src inhibitor (3,4-methylenedioxy-β-nitrostyrene) significantly reduced p-Src concentrations (Figure S2C-D). Thus, GIPR-mediated inhibition of Src may occur by direct interaction with the receptor, components of its signaling pathway, or by crosstalk with other membrane receptors that recruit Src family kinases (*29*).

As GIP reduced p-Src and bone resorption, we predicted that actin ring formation may be impaired by GIP. Bone slices were dyed with rhodamine and F-actin was labelled with SiR-actin, then time-lapse recordings captured by confocal microscopy. In osteoclasts exposed to GIP, the median time for initiation of actin ring formation leading to bone resorption was 26.2 hours, which was significantly longer than for vehicle-treated cells (10.9 hours, Figure 2E-F, Movie S1). Thus, GIP dependent suppression of bone resorption may be mediated by reduced p-Src signaling and delayed actin ring formation.

### GIP reduces PI3K-Akt signaling in osteoclasts

PI3K has been described to promote bone resorption by mechanisms including c-Src recruitment of PI3K (*30*) and activation of the serine-threonine kinase, Akt (*31*), while the PI3K-specific inhibitor wortmannin impairs actin ring and pit formation (*32*). To investigate PI3K signaling we examined Akt phosphorylation using an Akt1/2/3 (p-Akt) AlphaLISA assay, following 30 minutes exposure of osteoclasts to GIP or vehicle. P-Akt was significantly reduced by GIP, which was reversed by pre-treatment with GIP(3-30)NH_2_ (Figure 3A). These findings were replicated in osteoclast-osteoblast co-cultures (Figure S3A). Pre-treatment of cells with wortmannin phenocopies GIP actions on pAkt, while co-treatment with GIP and wortmannin has an additive effect on p-Akt production, which can be reversed by pre-treatment with the GIPR antagonist (Figure 3B). Thus, GIP acts at least in part on PI3K to reduce p-Akt generation in osteoclasts. However, as co-treatment with wortmannin and GIP could still further reduce p-Akt, GIP may activate additional pathways (Figure 3C).

**Figure 3.**
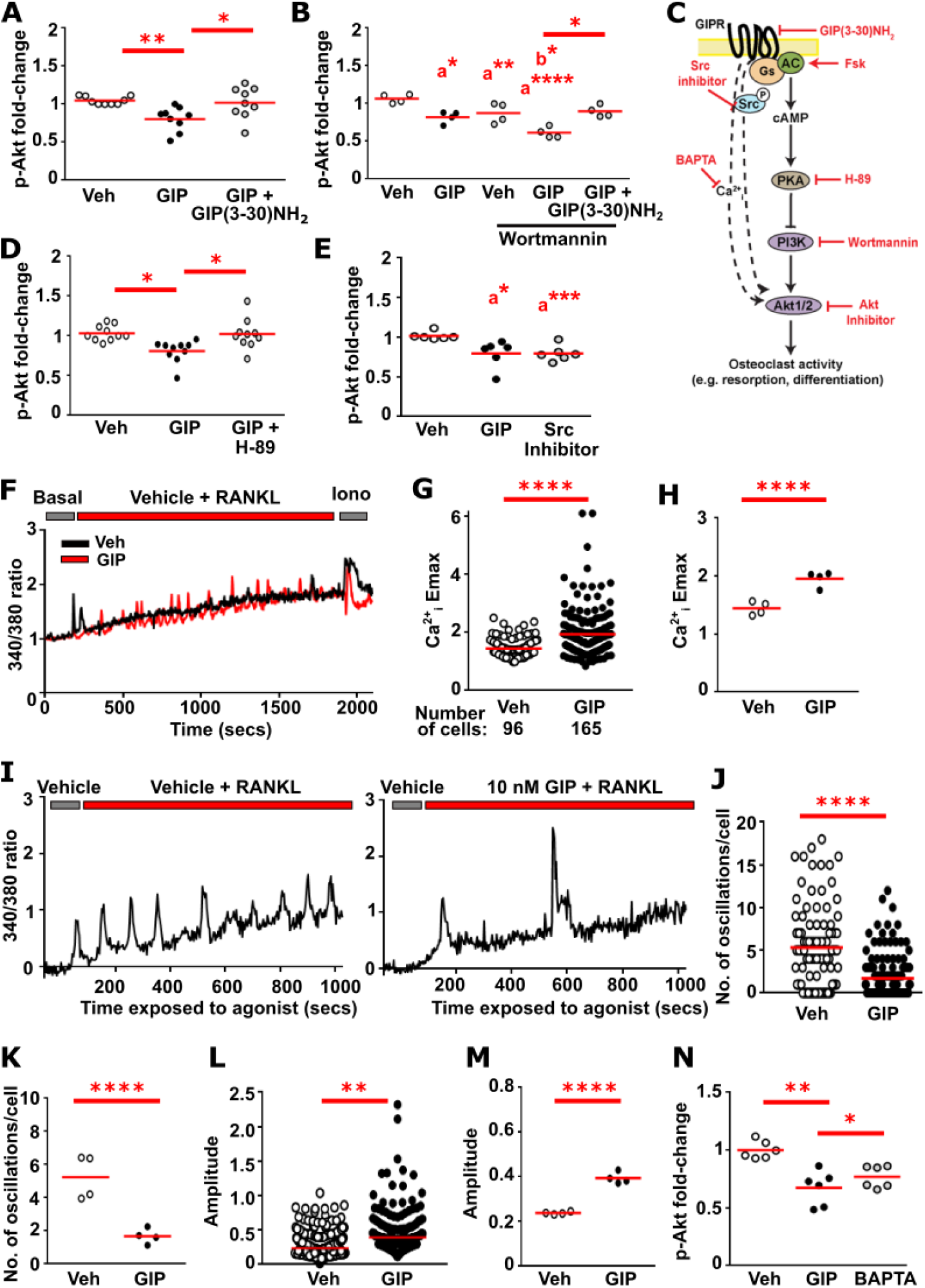
GIP reduces p-Akt signaling in human osteoclasts. (**A**) Quantification of phosphorylated Akt1/2/3 (p-Akt) generated in osteoclasts by vehicle (veh), GIP, or GIP with GIP(3-30)NH_2_ measured by AlphaLISA. (**B**) Effect of pre-treatment with the PI3K inhibitor wortmannin. (**C**) Cartoon showing GIPR signaling pathways that may act on Akt. Inhibitors/ activators are in red. (**D-E**) Effect of pre-treatment with (D) the PKA inhibitor H-89 and (E) a Src inhibitor on p-Akt responses. N=9 in A and B, N=10 in D and N=6 donors in E. Each dot represents one donor measured with median shown in red in panels A, B, D, E. (**F**) Normalized mean fluorescence intensity ratio (340/380nm) of Fura-2-AM calcium imaging in an individual cell exposed to vehicle or GIP. Data are normalized to the 340/380 ratio at 0 seconds for each cell. (**G-H**) Maximal Ca^2+^_I_ responses in all osteoclasts. G shows Emax of all cells measured. H shows average for each of the 4 donors. (**I**) Close-up of oscillations in a cell exposed to vehicle (left) and GIP (right). (**J-K**) Total number of Ca^2+^_i_ oscillations and (**L-M**) amplitude of oscillations from all cells measured. K and M show averages for each of the 4 donors. Data for individual donors is shown in Figure S3. (**N**) Effect of pre-treatment with the calcium chelator BAPTA on p-Akt. N=6 donors. p-Akt was normalized to GAPDH in AlphaLISA assays. Each dot represents one cell measured with median shown in red. ****p<0.0001, ***p<0.001, **p<0.01, *p<0.05. Comparisons to vehicle-treated cells are labelled as a and to GIP-treated cells as b. Statistical analyses were performed using: Kruskal-Wallis one-way ANOVA with Dunn’s multiple comparisons test for panel 3A, 3B, 3D, 3E, 3N; Mann-Whitney test for panels 3G, 3J, 3L; unpaired t-test for panels 3H, 3K and 3M.

Our studies had shown that GIP activates cAMP and c-Src pathways (Figure 1-2). To determine whether p-Akt is also reduced by GIP-mediated increases in cAMP, we pre-treated osteoclasts with the PKA inhibitor H-89 (Figure 3D). This prevented the GIP-induced reductions in p-Akt, indicating that GIP activates a cAMP-PKA pathway in osteoclasts. Moreover, exposure of osteoclasts to a Src inhibitor phenocopies the effect of GIP on osteoclasts (Figure 3E), indicating that the GIP-c-Src pathway could also contribute to reductions in p-Akt.

### GIP reduces calcium oscillations in osteoclasts to impair p-Akt generation

Previous studies in mouse osteoclast-like cell-lines showed exposure to a stable GIP analogue for 48 hours reduces RANKL-mediated Ca^2+^_i_ signaling (*11*), but did not investigate acute GIP exposure. To test this, we performed single-cell microfluorimetry with the calcium-indicating dye Fura-2. Following baseline readings for 180 seconds, osteoclasts were exposed to GIP or vehicle, before ionomycin addition in the final 180 seconds, which increases Ca^2+^ concentrations and serves as a positive control to determine cell responsiveness. Quantification of the 340/380 ratio showed that GIP induced large initial Ca^2+^_i_ responses and a greater maximal stimulatory response (Emax), than observed in vehicle-treated osteoclasts (Figure 3F-H, Figure S3). However, previous studies have shown that the frequency of calcium oscillations, rather than the maximal Ca^2+^_i_ response are important for governing the efficiency and specificity of gene expression in T lymphocytes (*33*), and for activation of NFATc1 and calcium/calmodulin-dependent protein kinase IV that are critical for osteoclast differentiation and resorption (*23, 34*). We therefore examined the frequency of Ca^2+^_i_ oscillations. Osteoclasts exposed to vehicle had frequent oscillations of similar amplitude, whereas osteoclasts treated with GIP had significantly fewer oscillations per cell (Figure 3I-K, Figure S3). The amplitude of these oscillations was significantly higher with GIP, consistent with elevated Emax (Figure 3L-M, Figure S3). Thus, GIP reduces Ca^2+^_i_ oscillation frequency, which is likely to reduce calcium-mediated gene transcription.

As previous studies have shown that p-Akt pathways can be stimulated by elevations in Ca^2+^_i_ (*35*), we hypothesized that reductions in Ca^2+^_i_ oscillations could contribute to the observed GIP-induced reductions in p-Akt. Indeed, p-Akt was lower in osteoclasts pre-treated with the calcium chelator BAPTA, such that p-Akt responses were not significantly different to GIP-treated cells. However, BAPTA did not reduce p-Akt levels when compared to vehicle-treated cells (Figure 3N), providing further evidence that multiple signaling pathways may act to reduce p-Akt downstream of GIPR in human osteoclasts.

### GIP impairs NFATc1 nuclear translocation by reducing Ca^2+^_i_-Akt and elevating cAMP

Ca^2+^_i_ oscillations and Akt play a critical role in activation of NFATc1, which on nuclear translocation regulates expression of genes involved in osteoclast differentiation and function (*36*). We therefore hypothesized that GIP may reduce NFATc1 translocation by the Ca^2+^_i_-Akt and cAMP-PKA-Akt pathways. Mature osteoclasts were exposed to RANKL with either vehicle or GIP for 60 minutes and immunocytochemistry performed examining NFATc1 expression in osteoclast nuclear and cytoplasmic fractions. Osteoclasts exposed to vehicle or pre-treated with GIP(3-30)NH_2_ had significantly greater NFATc1 in the nuclear fraction, when compared to GIP-treated cells (Figure 4A-C, Figure S4).Pre-treatment of cells with H-89 prevented the GIP-induced impairment in NFATc1 nuclear translocation (Figure 4A-C, Figure S4) indicating that GIPR-cAMP-PKA contributes to NFATc1 translocation. Co-treatment of cells with GIP and an Akt1/2 inhibitor further inhibited NFATc1 translocation, indicating that GIP-mediated reductions in Akt1/2 may also contribute to this pathway (Figure 4A-D, Figure S4).

**Figure 4.**
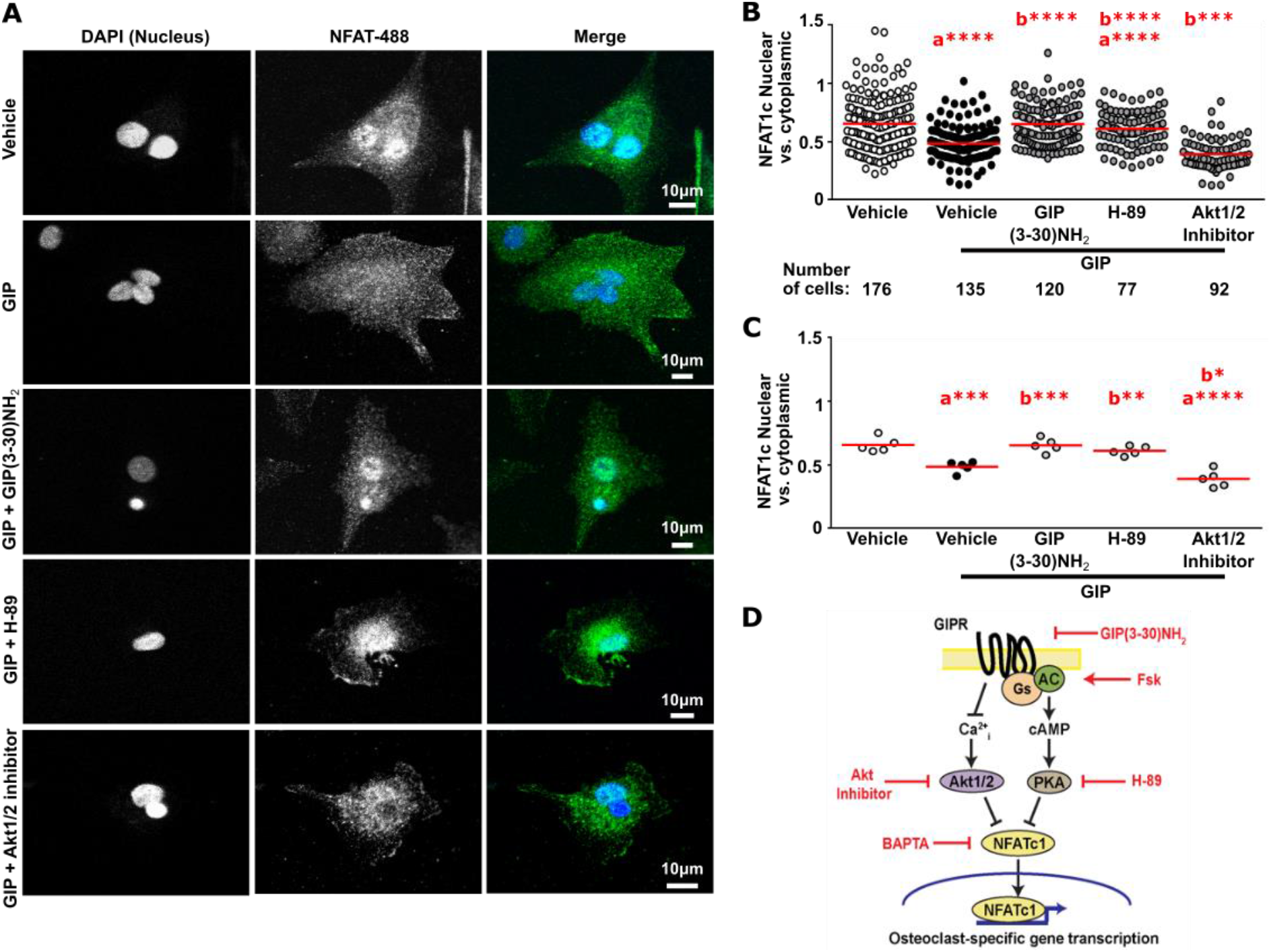
GIP reduces NFATc1 nuclear translocation in human osteoclasts. (**A**) Representative images of NFATc1 in osteoclasts exposed to vehicle or GIP +/- inhibitors of signaling. DAPI was used to label nuclei, and AlexaFluor488 to fluorescently label NFATc1. (**B-C**) NFATc1 nuclear and cytoplasmic ratios in osteoclasts from N=5 donors. B shows ratios for all cells measured. Each dot represents one donor measured with median shown in red in panel C. Data for individual donors is shown in Figure S4. Comparisons to vehicle-treated cells are labelled as a, and to GIP-treated cells as b. ****p<0.0001, ***p<0.001, **p<0.01, *p<0.05. (**D**) Cartoon showing GIPR signaling pathways that converge on NFATc1 activation. Inhibitors/ activators are in red. Statistical analyses were performed by: one-way ANOVA with Dunnett’s multiple comparisons test for 4B; and Kruskal-Wallis one-way ANOVA with Dunn’s multiple comparisons test for panel 4C.

### GIP reduces p38 signaling via Ca^2+^_i_-Akt pathways in osteoclasts

RANKL binding to RANK induces trimerization of the TNF receptor associated factor 6, leading to activation of the mitogen-activated protein kinase p38α and NFκB signaling pathways, resulting in enhanced osteoclastogenesis (*37*). p38 is activated by Src, can promote NFATc1 phosphorylation, interacts with Akt pathways, and mice with inducible deletion of p38α had reduced osteoclast numbers and decreased bone resorption (*38, 39*). To establish whether GIP affects p38 phosphorylation (p-p38), western blot analyses were performed of whole cell protein lysates from mature osteoclasts exposed to vehicle or GIP for 30 minutes. GIP produced significantly less p-p38 than osteoclasts exposed to vehicle or those pre-incubated with GIP(3-30)NH_2_ (Figure 5A). Therefore, p-p38 is impaired by GIPR activation.

**Figure 5.**
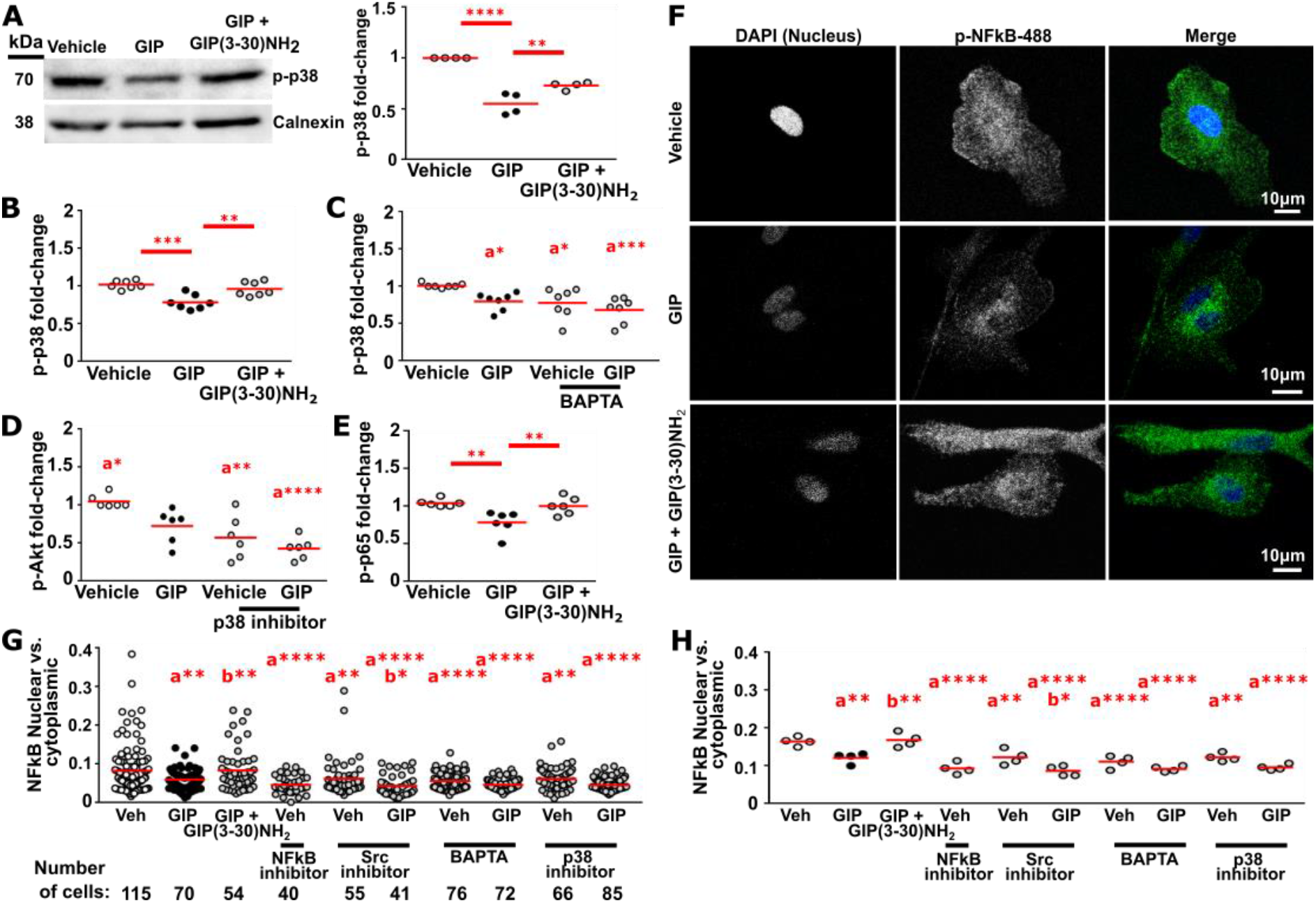
GIP reduces p38 and NFκB signaling in human osteoclasts. (**A**) Western blot analysis of phosphorylated p38 (p-p38) in osteoclasts exposed to vehicle or GIP for 30 minutes. (Right) Densitometry analysis of p-p38 relative to the calnexin loading control from 4 blots, each representing an independent donor. (**B**) Quantification of p-p38 measured by AlphaLISA. (**C**) Effect of pre-treatment with the calcium chelator BAPTA on p-p38 responses. (**D**) Effect of p38 inhibition on p-Akt in vehicle and GIP treated osteoclasts. (**E**) Quantification of phosphorylated p65 (p-p65), an NFκB subunit, in osteoclasts exposed to vehicle or GIP for 30 minutes measured by AlphaLISA. N=7 in B, N=6 donors in C-E. Data was normalized to GAPDH in AlphaLISA assays. (**F**) Representative images of p-NFκB in osteoclasts exposed to vehicle or GIP ± GIP(3-30)NH_2_ for 60 minutes. DAPI was used to label nuclei and AlexaFluor488 to fluorescently label p-NFκB. (**G-H**) p-NFκB nuclear and cytoplasmic ratios in osteoclasts from N=4 donors. G shows ratios for all cells measured. H shows average for each of the 4 donors. Data for individual donors is shown in Figure S5. Comparisons to vehicle-treated cells are labelled as a and to GIP-treated cells as b. ****p<0.0001, ***p<0.001, **p<0.01, *p<0.05. Each dot represents one donor measured with mean or median shown in red in panels A, B, C, D, E, H. Statistical analyses were performed by: one-way ANOVA with Tukey’s multiple comparisons tests for panels 5A and 5B, with Dunnett’s test for panels 5D and 5G, and with Holm-Sidak’s test for panel 5H. Kruskal-Wallis one-way ANOVA with Dunn’s test was used for panels 5D and 5G.

To determine the signaling pathways involved in GIP-mediated reductions in p-p38, AlphaLISA assays were performed with pathway inhibitors. These confirmed that p-p38 was decreased by GIP in osteoclast monocultures and osteoclast-osteoblast co-cultures and that pre-treatment with GIP(3-30)NH_2_ prevented this (Figures 5B and S5A). Pre-treatment with H-89, forskolin or a Src inhibitor had no effect on p-p38 responses (Figure S5B-D), indicating that the cAMP-PKA and Src pathways are unlikely to regulate p-p38. In contrast, pre-treatment of osteoclasts with the calcium chelator BAPTA reduced p-p38 in vehicle-treated cells, and combined GIP and BAPTA treatment had similar effects to GIP (Figure 5C). Finally, the effect of inhibition of the pAkt pathway on p-p38 was investigated. This showed that pre-treatment of osteoclasts with an Akt1/2 inhibitor had no effect on p-p38 (Figure S5E). However, inhibition of p38 impaired the generation of p-Akt (Figure 5D). Thus, GIP-induced reductions in p-p38 contributes to the inhibition of p-Akt, which may involve generation of Ca^2+^_i_ in human osteoclasts.

### GIP reduces NFκB signaling in osteoclasts

The transcription factor NFκB plays a critical role in osteoclast differentiation and function (*40*) and can be stimulated by PI3K-Akt and p38 (*41*), which we have shown are reduced by GIP (Figures 3-5). We therefore assessed phosphorylation of the p65 NFκB subunit (p-p65) by AlphaLISA. This demonstrated reduced p-p65 in GIP-treated osteoclasts and osteoclast-osteoblast co-cultures when compared to vehicle-treated cells, which was reversed by pre-treatment with GIP(3-30)NH_2_ (Figures 5E and S5F).

Following p65 activation, p65 translocates to the nucleus where it regulates transcription of genes that are essential for osteoclast differentiation and function. To assess NFκB nuclear translocation, mature osteoclasts were exposed to vehicle or GIP for 60 minutes and immunocytochemistry performed using a p-p65 antibody. Quantification of the nuclear vs. cytoplasmic p-p65 fractions showed that GIP reduced the amount of p-p65 in nuclear fractions to similar concentrations to that observed in cells exposed to an inhibitor of NFκB nuclear translocation (Figures 5F-H and S5). Therefore, GIP impairs the phosphorylation of NFκB subunits and reduces their nuclear translocation.

To decipher the mechanism by which GIP inhibits NFκB signaling in osteoclasts we pre-incubated osteoclasts with inhibitors of several signaling pathways. Exposure of osteoclasts to H-89 or forskolin had no effect on vehicle or GIP induced NFκB nuclear translocation (Figure S5). Pre-treatment with inhibitors of Src, Ca^2+^_I_, and p38 reduced NFκB nuclear translocation when compared to vehicle treated cells, indicating these pathways activate NFκB signalling in human osteoclasts (Figures 5G-H and S5). Combined treatments with GIP and inhibitors of Ca^2+^_I_ and p38 phenocopied the GIP effects (i.e. they did not have an additive effect), indicating that Ca^2+^_I_ and p38 may act downstream of GIP to inhibit NFκB signaling pathways (Figures 5G-H and S5).

### GIP reduces expression of genes involved in osteoclast activity and apoptosis

Our studies have shown that GIP reduces nuclear translocation of NFATc1 and NFκB, two major transcription factors regulating osteoclast gene expression (*36, 42, 43*) (Figure 4-5). We therefore hypothesized that GIP would affect osteoclast gene expression and performed RNA-seq analysis on mature osteoclasts exposed to vehicle or GIP for 4 hours. Analyses showed that there were 911 differentially expressed genes between the two groups (Figure 6A). Gene ontology analysis showed enrichment of genes involved in bone resorption, pH regulation, ATP production, apoptosis and the cell cycle (Figure 6B), correlating with osteoclast activity, and PI3K signaling, consistent with signaling assay data (Figure 6C). More than 40 genes were involved in lysosome and osteoclast function (Figure 6D). Lysosomes are required to generate the acidic environment of the lacuna, and to synthesize and secrete enzymes involved in bone extracellular matrix degradation. Among the downregulated genes we found those encoding for TRAP, cathepsin K and the calcitonin receptor, which are known to be regulated by NFATc1. Using the ISMARA algorithm (*44*) we quantified the impact of NFATc1 activity on global gene expression, which robustly increases during osteoclastogenesis, and decreases upon acute treatment with GIP in mature osteoclasts (Figure 6E). Genes predicted at higher confidence to be regulated by NFATc1, showed increasing specificity of osteoclast activity (Figure 6F) and higher susceptibility to be repressed by acute GIP treatment (Figure 6G), which was comparable across the 8 donors (Figure 6H).

**Figure 6.**
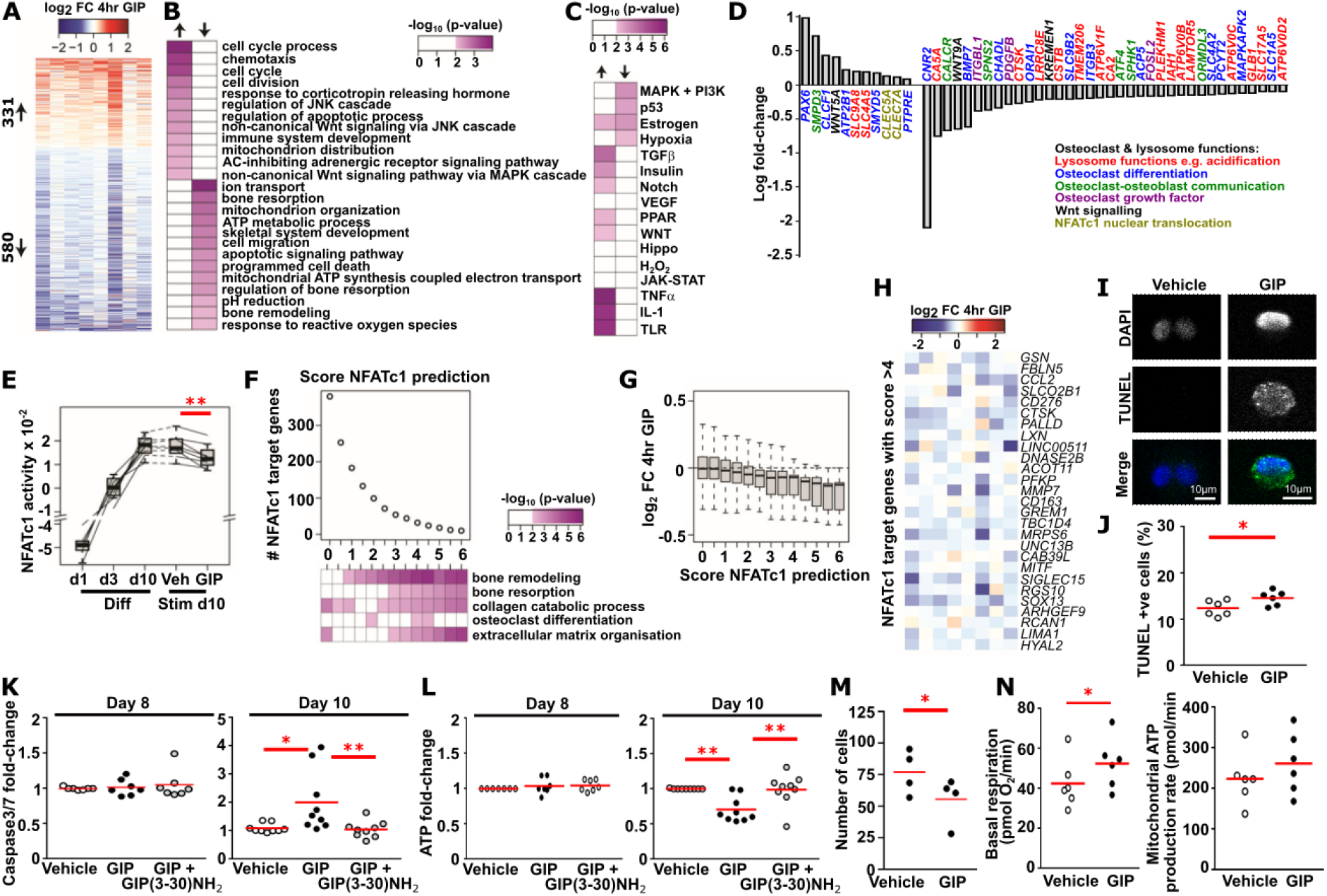
GIP reduces expression of genes involved in osteoclast function and induces apoptosis. (**A**) Heat map showing log fold changes (FC) for genes with significant changes (p-value <0.05) in expression upon 4 hours of GIP treatment in mature osteoclasts. N=8 donors, DEseq2 p-values modeled for donor and treatment. (**B**) Heat map showing enrichment for gene ontology terms among the up- and downregulated genes. (**C**) Heat map showing enrichment for targets of the indicated signaling pathways (SPEED2) among the up- and downregulated genes. (**D**) Log fold-change of significantly differentially expressed genes with known roles in osteoclast activities, comparing mature osteoclasts exposed to GIP for 4 hours to vehicle-treated cells. (**E**) ISMARA based motif activity of NFATc1 during osteoclast differentiation and in response to 4 hours GIP treatment in mature osteoclasts. (**F**) Heat map showing the enrichment of selected osteoclast related terms among NFATc1 targets with increasing confidence. (**G**) Box plot quantifying log fold-changes in gene expression of ISMARA based NFATc1 targets that were grouped according to prediction score upon 4 hours of GIP treatment. (**H**) Heat map showing the log fold changes for predicted NFATc1 target genes (ISMARA score above 4) in expression upon 4 hours of GIP treatment. (**I**) Representative images and (**J**) quantification of the percentage of TUNEL-FITC positive (+ve) osteoclasts following 3 days of GIP exposure. DAPI was used to stain nuclei. N=6 donors. (**K**) Caspase3/7 activity and (**L**) intracellular ATP levels, measured by CaspaseGlo and CellTiterGlo, respectively, in immature (day 8) and mature (day 10) osteoclasts exposed to vehicle, GIP, or GIP with GIP(3-30)NH_2_ for 3 days. N=7 for day 8, N=9 donors for day 10. (**M**) Total number of osteoclasts per bone slice following vehicle or GIP exposure. N=4 donors. (**N**) Basal respiration and mitochondrial ATP production rate in osteoclasts exposed to vehicle and GIP. N=6 donors. **p<0.01, *p<0.05. Each dot represents one donor measured, with mean or median shown in red in panels K-N. Statistical analyses were performed by: unpaired t-test for panel J; Kruskal-Wallis one-way ANOVA with Dunn’s test for panels 6K and 6L; and paired t-test for panels 6M and 6N.

Many genes involved in apoptosis or mitochondrial function were differentially expressed in osteoclasts exposed to GIP (Figure 6B). We therefore sought to determine whether GIP affects apoptosis in human osteoclasts. Initially, terminal deoxynucleotidyl transferase dUTP nick end labeling (TUNEL) staining was performed. Mature osteoclasts exposed to GIP for 3 days had a significantly greater number of TUNEL-positive cells when compared to vehicle-treated cells (Figure 6I-J). In line with these findings, caspase-3/7 apoptosis assays demonstrated increased apoptosis in mature osteoclasts exposed to GIP, but not in cells pre-treated with GIP(3-30)NH_2_ (Figure 6K). GIP did not increase apoptosis in immature osteoclasts (Figure 6K).

Mitochondrial function is critical for osteoclast differentiation and bone resorptive activity (*45*). To determine whether osteoclasts were less metabolically active, we measured intracellular ATP. While levels were unaffected by GIP in immature osteoclasts, intracellular ATP concentrations were lower in GIP-treated mature osteoclasts, indicating these cells may be less metabolically active (Figure 6L). ATP levels may be decreased by a reduction in the total number of osteoclasts or due to changes in osteoclast energy metabolism. Consistent with our findings that GIP increases apoptosis, GIP reduced the total number of osteoclasts (Figure 6M). To determine whether alterations in osteoclast energy metabolism also affect ATP production, we performed bioenergetic profiling using extracellular flux assays. Acute treatment with vehicle or GIP (0.1, 1, 10, 100 nM) for 30 minutes showno differences in basal metabolism (Table S2). To evaluate if longer treatments with GIP are required to alter the mitochondrial function, we performed bioenergetic profiling in mature osteoclasts exposed to GIP for 3 days. A small but significant increase in basal respiration was observed in cells exposed to GIP (1.23 fold-change)), with a tendency to increased mitochondrial and total ATP production rate (Table S3, Figure 6N). No other significant differences were observed. Thus, it is likely that GIP does not have a major effect on osteoclast bioenergetics, and that the reduced ATP levels could be explained by fewer cells and/or number of nuclei per osteoclast caused by GIP treatment.

### GIP increases p-Akt in human osteoblasts via a cAMP-PKA pathway, leading to decreased cell apoptosis

GIPR expression has been shown in osteoblast-like cell-lines (*7, 9*), and we confirmed *GIPR* expression in mature primary human osteoblasts using qPCR (Figure S7A). To determine the functional relevance, we stimulated GIPR signaling in human osteoblasts, cAMP accumulation was measured at 4 time points (15, 30, 60, 120 minutes). This showed a significant elevation in cAMP in cells treated with GIP for 15 and 30 minutes when compared to vehicle-treated cells (Figure S7B). Pre-treatment with GIP(3-30)NH_2_ prevented GIP-mediated increases in cAMP, indicating direct stimulation of GIPR on osteoblasts (Figure 7A). Therefore, GIP can directly activate GIPR-mediated signaling in human osteoblasts.

**Figure 7.**
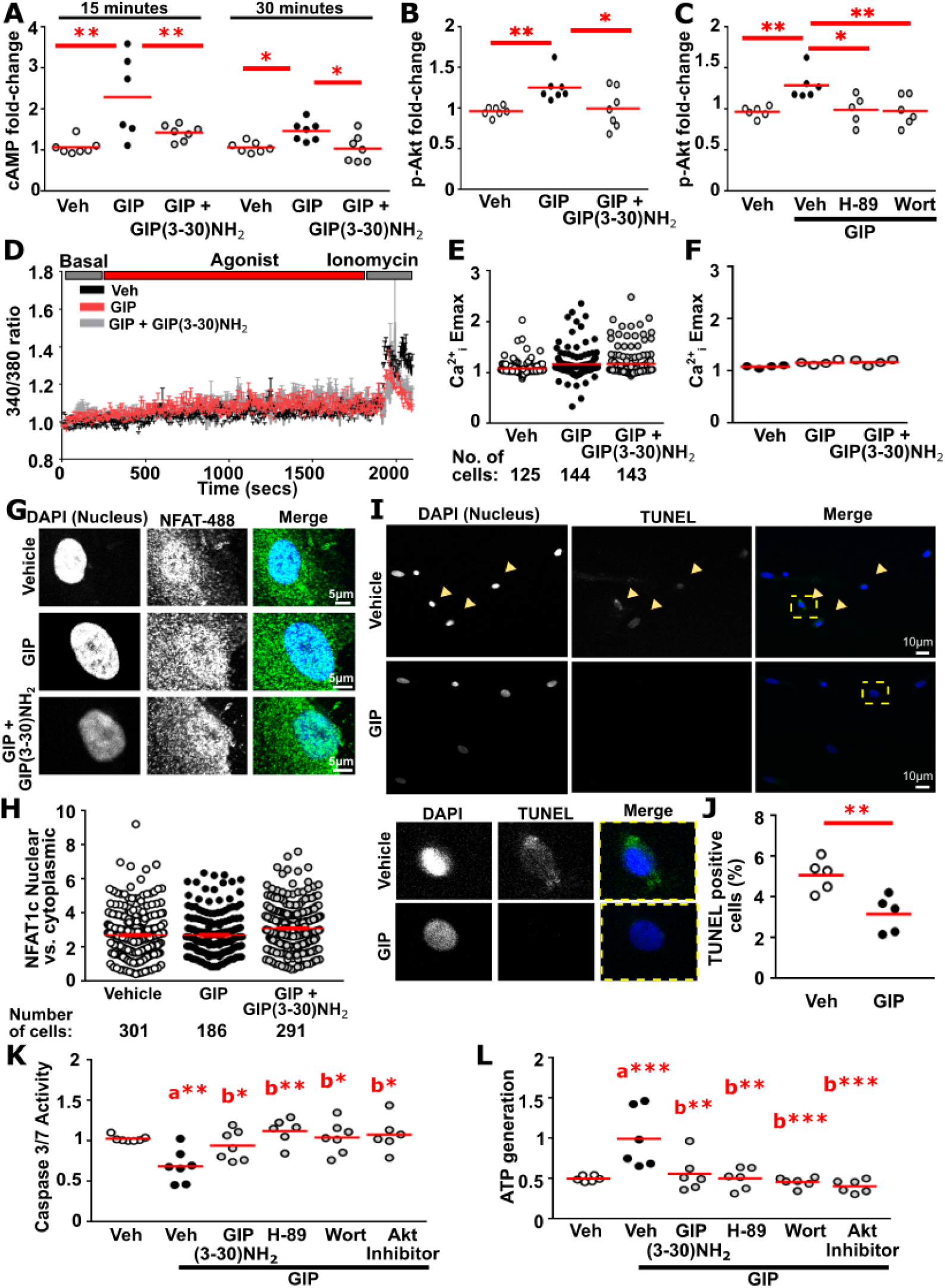
GIP increases cAMP and pAkt signaling, and reduces apoptosis in human osteoblasts. (**A**) Quantification of cAMP in osteoblasts exposed to vehicle (veh), GIP or GIP+GIP(3-30)NH_2_ measured by LANCE assays. N=7 donors. (**B-C**) Quantification of phosphorylated Akt1/2/3 (p-Akt) in osteoblasts exposed to vehicle or GIP for 30 minutes measured by AlphaLISA. Cells were pre-treated with vehicle and (B) GIP(3-30)NH_2_, (C) the PKA inhibitor H-89 and PI3K inhibitor wortmannin. N=7 for B, N=6 donors for C. p-Akt concentrations were normalized to GAPDH in AlphaLISA assays. (**D**) Normalized mean fluorescence intensity ratio of Fura-2-AM. Data are normalized to the 340/380 ratio at 0 seconds for each cell. Data shows mean+SEM. N=4 donors. (**E-F**) Quantification of the maximal Ca^2+^_I_ responses from data in panel D. E shows Emax of all cells measured. F shows average for each of the 4 donors. (**G**) Representative images of NFATc1 and (**H**) quantification of nuclear and cytoplasmic ratios in osteoblasts exposed to vehicle or GIP for 60 minutes. DAPI was used to label nuclei and AlexaFluor488 to fluorescently label NFATc1. N=5 donors. (**I**) Representative images of osteoblasts stained with TUNEL to detect apoptotic cells and DAPI to detect nuclei, with zoomed images of cells indicated by a yellow box shown below. (**J**) Quantification of the percentage of TUNEL-FITC positive cells. N=5 donors (**K**) Caspase-3/7 activity and (**L**) ATP generation, measured by CaspaseGlo and CellTiterGlo, respectively, in osteoblasts exposed to vehicle, GIP or GIP +/- inhibitors of signaling. N=6 donors. Comparisons to vehicle-treated cells are labelled as a, and to GIP-treated cells labelled as b, in panels J and K. ***p<0.001, **p<0.01, *p<0.05. Each dot represents one donor measured with mean or median shown in red in panels A-E, G-H. Statistical analyses were performed by: Kruskal-Wallis one-way ANOVA with Dunn’s test for panels 7A, 7B, 7E, 7F, 7H, 7K; one-way ANOVA with Dunnett’s multiple comparisons test for 7C and 7L; and unpaired t-test for 7J and 7M.

Previous studies showed that Akt1 and the RUNX2 transcription factor enhance osteoblast differentiation (*46*). We therefore assessed whether GIP influenced p-Akt signaling in osteoblast monocultures using AlphaLISA assays. Exposure of osteoblasts to GIP increased p-Akt concentrations, which was prevented in cells pre-treated with GIP(3-30)NH_2_ (Figure 7B). Additionally, pre-treatment with H-89 or wortmannin inhibited GIP-induced increases in p-Akt, demonstrating PKA and PI3K dependent p-Akt activation in response to GIP in osteoblasts (Figure 7C). In contrast to osteoclasts, Ca^2+^_i_ signaling and NFATc1 nuclear translocation were unaffected by GIP in osteoblasts (Figure 7D-H, Figure S7C-D). Studies have shown that GIP activates p38 signaling in mouse osteoblast-like cells (*47*). However, no difference was observed in p-p38 in human osteoblasts exposed to vehicle or GIP assessed by AlphaLISA (Figure S7E). Finally, we assessed apoptosis in osteoblasts as Akt signaling can promote cell survival pathways. Staining revealed significantly fewer TUNEL positive cells in osteoblasts exposed to GIP, when compared to vehicle (Figure 7I-J), and caspase3/7 was reduced by GIP (Figure 7K). Pre-treatment with GIP(3-30)NH_2_, H-89, wortmannin or an Akt1/2 inhibitor prevented the GIP-induced reduction in caspase3/7 activation (Figure 7K), indicating that GIPR activation of cAMP-PKA and PI3K-Akt signaling pathways are involved in GIP-mediated reductions in osteoblast apoptosis. Consistent with this, GIP increased ATP generation in osteoblasts, which was reduced by GIP(3-30)NH_2_, H-89, wortmannin and an Akt1/2 inhibitor (Figure 7L).

## Discussion

Our studies demonstrated that GIPR is robustly expressed on primary human osteoclasts, with increased expression during the differentiation process, and that GIP acts upon multiple signaling pathways to reduce osteoclast differentiation and bone resorption (Figure S7). GIPR is also expressed on proliferating mature human osteoblasts and GIP acts upon a cAMP-Akt pathway to improve osteoblast survival. Previous studies have shown that endogenous GIP contributes to the postprandial suppression of bone resorption by up to 25% in humans (*48*), and that GIP abruptly decreases circulating biomarkers of bone resorption (*9, 16–18*). In addition, targeting whole-body GIPR, improved bone strength in mice with ovariectomy-induced bone loss (*47*). However, these studies did not determine whether changes in bone turnover were due to direct effects on GIPR on bone cells, or indirect effects via extra-skeletal GIPR. A major advantage of our studies is in the use of the GIPR antagonist, GIP(3-30)NH_2_, in all assays, to demonstrate direct effects of GIP on osteoclast and osteoblast expressed GIPR in human primary cell cultures. Other gut-derived hormones, including GLP-1 and GLP-2, have been described to affect bone (*17, 49*), and adoption of a similar strategy to investigate bone cell signaling and activity with specific antagonists would help determine whether these hormones induce direct effects on human bone cells.

We showed that GIP stimulates the canonical GIPR cAMP pathway, and that this pathway contributes to reduced osteoclast activity. This is consistent with previous studies that showed forskolin, which elevates cAMP by activating adenylate cyclase, impairs bone resorption (*50*). Previous studies conducted in mouse cell-lines did not detect GIP-induced changes in cAMP (*11*), and there are several possible reasons for these differences. Our studies investigated primary human osteoclasts, and it is possible that different signaling pathways exist between each species, or between primary and immortalized cells. However, we were able to verify other GIP-induced signaling effects (i.e., inhibition of Ca^2+^_I_ and impaired NFATc1 nuclear translocation) observed in these mouse cell-lines. Alternatively, experimental differences could explain why cAMP induction was not previously observed. We used a highly sensitive LANCE cAMP assay, which unlike the FRET probe used in the mouse cell-lines does not require transfection. Due to these differences, it will be important to investigate the effects of newly designed GIPR analogues in primary human bone cells, rather than mouse cell-lines, to assess possible effects on osteoclast activity.

Our examination of multiple signal readouts demonstrated that GIP regulates several signaling pathways in human osteoclasts and that these have distinct effects on gene expression. Thus, NFATc1 nuclear translocation is regulated by cAMP-PKA and Ca^2+^_i_ signaling pathways, whereas NFκB is regulated by Src and Ca^2+^_i_-p38-Akt, but not by cAMP-PKA. This could explain why GIP affects multiple aspects of osteoclast function including delaying actin ring formation, altering expression of genes involved in osteoclast development and function, increasing apoptosis, and ultimately impairing osteoclast differentiation, survival and bone resorption, which together explain the potent reduction in resorption *in vitro* in our studies, and GIP-induced reductions in bone resorption markers in human studies (*9, 16–18*). We showed that GIP-induced signaling events in human bone cells occur rapidly (phosphorylation events by 30 minutes, nuclear translocation by 60 minutes and gene expression by 4 hours), consistent with known acute signaling by GIPR (*51*). Previous studies had shown that NFATc1 nuclear translocation was impaired by chronic exposure (48 hours) to GIP (*11*); however, chronic GIPR activation desensitizes GIPR activity (*52*). Thus, our studies demonstrate that GIP-mediated effects on human osteoclasts are due to acute activation of the GIPR, rather than indirect effects of receptor desensitization.

Despite an enrichment in differentially expressed genes associated with metabolic pathways and oxidative phosphorylation, and significantly lower ATP concentrations in osteoclasts exposed to GIP, we did not identify any changes in ATP production. The most likely explanation for lower ATP concentrations is that GIP, as observed, reduces the total number of osteoclasts and the number of nuclei per osteoclast, possibly due to increased apoptosis. However, the differences identified between total ATP concentrations and ATP production in GIP stimulated osteoclasts could reflect more complex dynamics between intracellular and extracellular ATP that have previously been shown to regulate osteoclast survival and bone resorption (*53*).

GIP had a more pronounced effect on osteoclasts than osteoblasts in our studies, presumably due to higher osteoclastic GIPR expression. However, osteoblast GIPR remains functional, resulting in activation in signaling pathways, reduced apoptosis and improved survival of osteoblasts, which may contribute to the transient and small increase in bone formation markers observed in some human studies (*18, 19, 49, 54*). This indicates that GIP alters bone remodeling, causing a decrease in bone resorption while maintaining bone formation, and the increased survival combined with reduced apoptosis could improve osteoblast function. Current anti-resorptive treatments for osteoporosis, such as bisphosphonates, inhibit osteoclast activity and through coupling mechanisms also osteoblast activity leading to lower bone remodeling, which may compromise skeletal integrity with long-term exposure (*55*). Thus, targeting GIPR could have advantages over bisphosphonates as our studies demonstrates that GIP inhibits osteoclast activity, but increases osteoblast survival, which could lead to larger gains in bone mass and possibly lower fracture risk. Importantly, it remains to be studied in clinical studies if the observed changes in bone cell activity are maintained with long-term GIPR agonism. Drugs targeting the GIPR are currently being investigated as treatments for metabolic conditions such as type 2 diabetes (*56–58*). Our study indicates that these may have advantageous effects on bone remodeling. Therefore, it is important to investigate the effect of these drugs on circulating bone turnover markers and bone resorption and formation, to establish whether chronic GIPR activation uncouples bone remodeling, which could emerge as a novel approach to treating metabolic bone diseases, such as osteoporosis and skeletal fragility as observed in type 2 diabetes (*59*).

In conclusion, our studies have shown that GIP acts via multiple signaling pathways in osteoclasts, to reduce differentiation and bone resorption, and activates GIPR in osteoblasts to improve cell survival. Thus, stable GIP analogues may reduce bone resorption without impairing bone formation, supporting studies of long-term effects of GIP on bone.

## Materials and Methods

### Compounds

Compounds used in these studies were used at the following concentrations and purchased from companies as stated: Akt1/2 inhibitor (100nM, Sigma Catalog A6730), BAPTA (10μM, Sigma), c-Src inhibitor 3,4-methylenedioxy-β-nitrostyrene (20nM, SantaCruz), Forskolin (10μM, Tocris), GFX (2μM, GF-109203X, GFX), human GIP (0.1, 1, 10 and 100nM, Bachem), human GIP(3-30)NH_2_ (10µM, Caslo), H-89 dihydrochloride (10μM, Tocris), NFκB nuclear translocation inhibitor (100μg/ml, SantaCruz, Catalog sc-3060), p-38 inhibitor, PH-797804 (25nM, Tocris), Wortmannin (1mM, Tocris). All studies were performed in the absence of DPP4 inhibitors (i.e. with endogenous DPP4) to better replicate the native environment of *in situ* osteoclasts and osteoblasts.

### Generation of human osteoclasts and osteoblasts

Primary human osteoclasts were differentiated from human CD14^+^ monocytes isolated from buffy coats obtained from anonymous blood donations as previously described (*22*). Monocytes were seeded in T75 cell culture flasks (5 x 10^6^ cells/flask) in α-minimal essential medium (αMEM, Gibco) with 10% fetal bovine serum (FBS, Gibco or Sigma), 1% penicillin/streptomycin (Gibco) and stimulated with 25 ng/mL macrophage colony-stimulating factor (MCSF, R&D Systems). After two days, media was replaced with fresh media containing MCSF and receptor activator of nuclear κB (RANKL) (25 ng/mL each, R&D Systems), and media refreshed every 2-3 days thereafter until a total of seven days stimulation with RANKL. Microscopic validation of the presence of multinucleated cells was performed before cells were used for experiments. All cells were maintained at 37°C, 5% CO_2_.

Primary human osteoblast lineage cells were obtained from bone specimens from patients receiving hip replacement surgery due to osteoarthritis, as previously described (*60*). The bone was cut into small pieces (∼5 mm in diameter) and cleaned in PBS. Five bone pieces were placed in each well of a 12-well plate containing Dulbecco’s modified Eagle medium (DMEM, Gibco), 10% FBS (Sigma), 2nM L-glutamine (Gibco), 50 µg/mL ascorbic acid (Sigma), 10 mM β-glycerophosphate (Sigma) and 10^-8^ M dexamethasone (Sigma). To prevent the bone pieces from moving, a metal grid was placed on top of the bone slices, and samples incubated for fourteen days, with a single media change at day seven. On day fourteen, metal grids were removed, cells expanded to larger flasks, with media changes twice weekly until the osteoblast lineage cells reached near confluency after a total of ∼35 days.

All cells were maintained at 37°C, 5% CO_2_. Clinical investigations were conducted in line with principles expressed in the Declaration of Helsinki and informed consent was obtained from all donors. Approval for collection of samples for cell culture and *in situ* hybridization were obtained from local ethics committees in Denmark (S-2011-0114 & S-20120193) and the UK (ERN_14-0446).

### *In situ* hybridization

Formalin-fixed, decalcified, and paraffin-embedded bone specimens were obtained from the proximal femur of eight human adolescent patients during corrective surgery for coxa valga, as previously described (*61*). Paraffin sections (3.5-μm-thick) were subjected to *in situ* hybridization using an enhanced version of the RNAScope 2.5 high-definition procedure (310035, ACD Bioscience). Sections were rehydrated, deparaffinized, and pretreated, as previously described (*62*), using protease instead of pepsin, and hybridized overnight at 40°C with 20-ZZ-pair probes (477541,ACD Bioscience) directed against the 384-1553 region of human GIPR mRNA (NM_000164.2) diluted 1:1 in probe diluent (449819, ACD Bioscience). Negative controls hybridized with probe diluent only were included for each donor. Amplification was conducted according to the instructions provided by the manufacturer. Horseradish peroxidase was further enhanced with digoxigenin (DIG)-labeled tyramide (NEL748001KT, PerkinElmer), which was labeled with AP-conjugated sheep anti-DIG FAB fragments (11093274910, Roche) and visualized with Liquid PermanentRed (Dako). Finally, the sections were counterstained with Mayer’s hematoxylin and mounted with Aqua-Mount. Data analysis was performed blinded.

### RNAseq

CD14^+^ monocytes were isolated from 8 different donors and seeded in T25 cell culture flasks (1.67×10^6^ cells/flask). Cells were differentiated as described above with 10nM GIP or vehicle added on day 10 for 4 hours before cells were harvested using TRIzol. RNA was purified using Econo Spin columns (Epoch Life Sciences). RNA-sequencing was performed according to manufacturer’s instructions (TruSeq 2, Illumina) using 2µg RNA for preparation of cDNA libraries. Sequencing reads were mapped to the human genome (GRCh38) using Spliced Transcripts Alignment to a Reference (STAR) software (*63*), and tag counts were summarized at the gene level using Hypergeometric Optimization of Motif EnRichment (HOMER) (*64*). Differential gene expression was analyzed using DESeq2 (*65*) taking both donor and stimulation effects into the model. Gene ontology analysis was performed using goseq (*66*) and pathway analysis was performed using the SPEED2 (doi: 10.1093/nar/gkaa236) database with a hypergeometric test or the Database for Annotation, Visualization and Integrated Discovery (DAVID) v6.8. Motif activity and target prediction analysis used the ISMARA algorithm (doi: 10.1101/gr.169508.113) by submitting raw sequencing data. Raw RNA-seq data and DESeq2 processed data that have been generated in this study are available under the GEO accession number GSE201100.

### Osteoclast resorption assays

Multinucleated osteoclasts were seeded on bovine cortical bone slices (0.4 mm thickness) (BoneSlices.com) in 96-well plates at a density of 50,000 cells per bone slice in αMEM with 10% FBS, 25 ng/mL MCSF and 25 ng/mL RANKL (n=6 technical replicates per donor). Cells were allowed to settle for 40 minutes, then vehicle or GIP (0.1, 1, 10 or 100 nM) added to the media. In studies with the GIPR antagonist, cells were pre-incubated with 10µM GIP(3-30)NH_2_ or vehicle (water) for 10 minutes before adding 10nM GIP. For osteoclast and osteoblast co-cultures, 50,000 osteoclasts per bone slice were seeded without FBS or RANKL on bovine bone slices allowed to settle for four hours. Osteoblasts were then seeded on top of the osteoclasts at a density of 12,500 cells per bone slice before adding agonist and antagonist as described above (*60*).

For monoculture and co-culture studies, cells were incubated for 72 hours, then media collected for use in subsequent studies, and the experiment terminated by adding 200µl demineralized water to each bone slice. Bone slices were scraped with a cotton swab, and stained with Toluidine Blue Solution (1% toluidine blue, 1% sodium borate) (Sigma-Aldrich) to visualize resorption excavations using a 100-point counting grid and counting performed using a 10x objective of a BX53 Olympus microscope (Olympus). Bone resorption was assessed blinded by calculating the total eroded surface percentage (%ES), as previously described (*67*).

### Quantification of the number of nuclei per osteoclast

CD14^+^ monocytes were isolated and seeded as described. After 2 days, cells were loosened with Accutase and seeded in 96-well plates (2.5 × 10^4^ cells per well). Cells were incubated with media containing 25 ng/ml MCSF and 25 ng/ml RANKL and either vehicle (water) or 10nM GIP (n=6 technical replicates, 4 donors). Media containing either vehicle or 10nM GIP was refreshed after 3 and 5 days. After 7 days of culture with vehicle or GIP, media was removed, and the wells were washed twice with PBS. Cells were then fixed with 3.7% formalin and methanol and stained with Giemsa and May-Grünwald (Merck), as previously described (*68*). The number of multinucleated osteoclasts and their number of nuclei were counted systematically in every second counting field using an Axiovert 200 microscope (Zeiss).

### Tartrate-resistant acid phosphatase 5b (TRAcP) activity analysis

TRAcP activity was measured in conditioned media from each of the bone resorption experiments. Conditioned media (10µl per well) was transferred in duplicates into a 96-well plate with 90µl TRAcP solution buffer (1M acetate, 0.5% Triton X-100, 1M NaCl, 10 mM EDTA (pH 5.5), 50 mM L-Ascorbic acid, 0.2 M disodium tartrate, 82 mM 4-nitrolphenylphosphate, all reagents from Sigma) and incubated at 37°C in the dark for 30 minutes. The reaction was stopped by adding 100 µl stop buffer (0.3M NaOH) and TRAcP activity was measured at absorbance 400 nm and 645 nm on a Synergy HT microplate reader (Biotek instruments).

### LANCE cAMP assays

Mature osteoclasts were plated on bone slices in 96-well plates at a density of 6,250 cells per well in osteoclast monocultures and co-cultures. In co-culture studies, osteoclasts were seeded in RANKL-free media and left to settle, before 1,500 osteoblasts per well were added. Cells were exposed to either vehicle (water) or 10nM GIP in stimulation buffer (1x Hanks Buffered Saline Solution, 0.1% BSA, 0.1% 3-isobutyl-1-methylxanthine (IBMX) (all Sigma), 0.5mM HEPES (Invitrogen)) for either 0, 15, 30, 60 or 120 minutes. LANCE cAMP assays were then performed following the manufacturer’s instructions. Immediately prior to plate reading, the contents of the wells were transferred to a white 96-well plate and HTRF readings made on a Pherastar FS (BMG Labtech) plate reader. The HTRF ratio (Em665/Em615) values were normalized to vehicle treated cells and expressed as 1/ to yield positive values. Graphs were generated using GraphPad Prism 7.

### AlphaLISA phosphorylation assays

Mature osteoclasts were plated on bone slices in 96-well plates at a density of 10,000 cells/ well in osteoclast monocultures and co-cultures. In co-culture studies, osteoclasts were seeded in RANKL-free media and left to settle, before adding 1,500 osteoblasts per well. Cells were exposed to either vehicle (water) or 10nM GIP in stimulation buffer for 30 minutes, then lysed in AlphaLISA buffer (PerkinElmer). For studies with inhibitors, cells were pre-treated with each compound for one hour at the concentrations indicated in the *Compounds* section. For studies with forskolin, comparisons were made to an ethanol vehicle, while DMSO was the vehicle for all other compounds.

AlphaLISA assay kits to detect GAPDH, as a measure of total protein, and phosphorylated CREB (Ser133), c-Src (Tyr419), Akt1/2/3 (Ser473), p65 NFkB (Ser536) and p38 (Thr180/Thr182) were purchased from PerkinElmer and performed according to manufacturer’s instructions. Assays were performed in white 384-well plates (Optiplates, PerkinElmer) and AlphaLISA readings made on a Pherastar FS (BMG Labtech) plate reader. Values for phosphorylated proteins were normalized to GAPDH values. Graphs were generated using GraphPad Prism 7.

### Time-lapse recording of osteoclasts (actin ring formation)

Time-lapse recording of osteoclast activity on bovine bone slices was performed, as previously described (*69*). In brief, bone slices with a thickness of 0.2mm (BoneSlices.com) were labelled with N-hydroxysuiccinimide ester-activated Rhodamine fluorescent dye (ThermoFisher). Mature osteoclasts were seeded at a density of 100,000 cells/bone slice in a 96-well plate. F-actin was labelled using 100nM SiR-actin (excitation at 652 nm; emission at 674 nm) and 10µM verapamil (both supplied by Spirochrome) and incubated for 4 hours at 37°C, 5% CO_2_. Subsequently, bone slices were inverted and placed in a Nunc Lab-Tek II chambered cover-glass (ThermoFisher Scientific) in medium containing M-CSF, RANKL, SiR-actin and verapamil with vehicle or 10nM GIP (n=2 technical replicates per donor, n=2 donors). Time-lapse images were made using an Olympus Fluoview FV10i microscope (Olympus Corporation) at 37°C, 5% CO_2_, with a 10× objective and a confocal aperture of 2.0 corresponding to a *z*-plane depth of 21.2 µm. Recordings were made for a period of 72 hours taking images every 21 min.

### Intracellular calcium imaging

Cells were plated in six-well plates at 100,000 cells per well on coverslips. Cells were left to settle then pre-incubated with media containing 100nM GIP(3-30)NH_2_ or vehicle control (DMSO) for one hour. Cells were loaded with 4μM Fura-2 AM (Molecular Probes) dissolved in DMSO with 0.03% pluronic acid (Sigma) for 30 mins and imaged on a Crest X-light spinning disk system coupled to a Ti-E base and 10x/0.4 air objective. Excitation was delivered at l = 340 nm and l = 385 nm using a FuraLED system, with emitted signals detected at l = 470–550 nm, as described (*70*). Cells were imaged in HBSS with 10mM HEPES (Gibco) at 37°C. Cells were imaged for 3 minutes (60 frames) in imaging buffer, then 10nM agonist (vehicle or GIP with RANKL) added and a further 660 frames imaged, prior to addition of the positive control 1μM ionomycin (Sigma) for 60 frames. Live sequential images were captured at 340nm and 380nm excitation, and emitted signals detected at l = 470–550nm using a FuraLED system. Data was captured using MetaMorph software (Molecular Devices) and calculations of 340/380 ratios performed in ImageJ. For each dataset, ratio values were normalized to the ratio in the first frame that was set at 1. Data was plotted in GraphPad Prism.

### NFATc1 and NFκB confocal microscopy

Cells were plated in six-well plates at 100,000 cells per well on coverslips. At least four hours later, cells were pre-incubated with inhibitors at concentrations described in *Compounds* for one hour. Vehicle (water) or 10nM GIP was added to cells for 60 minutes, then cells washed in PBS and fixed in 4% paraformaldehyde/PBS (SigmaAldrich). Cells were permeabilized in 1% Triton-X100/PBS (Thermo Scientific), and immunostained with either total NFATc1 or p-p65 primary antibodies (1:1000, Cell Signaling Technologies) and secondary antibody Alexa Fluor 488 (1:3000, Invitrogen). Cells were mounted in Prolong Gold Antifade reagent (Invitrogen), which counterstains cell nuclei with DAPI. Images were captured using a Zeiss LSM 780 confocal microscope with a Plan-Apochromat x63/1.2/water DIC objective. An argon laser (488 nm) was used to excite Alexa Fluor 488. The mean fluorescence intensity per pixel was measured in regions of interest within the nucleus and cytoplasm to calculate nuclear-to-cytoplasmic ratios using ImageJ (NIH). Each assessor was blinded to the culture conditions. Ratios were then plotted in GraphPad Prism 7. For studies with forskolin, comparisons were made to an ethanol vehicle, while DMSO was the vehicle for all other compounds.

### Western blot analysis

Mature osteoclasts were plated in 6-well plates at a density of 100,000 cells/well and allowed to settle for four hours. Cells were pre-incubated with media containing 100nM GIP(3-30)NH_2_ or vehicle control (DMSO) for one hour, prior to addition of 10nM GIP or vehicle (water) for 0 and 30 minutes. Western blots were then performed as previously described (*71*). Following incubation, cells were lysed in NP40 lysis buffer (50mM Tris HCl pH7.4, 1mM EDTA, 150mM NaCl, protease inhibitors (all purchased from Sigma)), lysates were resuspended in Laemmli buffer, boiled and separated on 12% sodium-dodecyl sulphate (SDS) polyacrylamide gel electrophoresis gels. Following transfer to polyvinylidene difluoride membrane (Amersham), blots were blocked in 5% BSA/TBS-t, then probed with the primary p-p38 (1:1000, Cell Signaling Technologies) and donkey-anti-rabbit antibody (1:3000, Invitrogen). Blots were visualized using the Immuno-Star WesternC kit (BioRad) on a BioRad Chemidoc XRS+ system. Following development, blots were stripped with Restore Western blot stripping buffer (Thermo Scientific), blocked in marvel/TBS-t and re-probed with a calnexin antibody (1:1000, Millipore).

### Apoptosis and cell viability (ATP) assays

Osteoclasts were plated in 96-well plates at 6,250 cells/well on day 8 or day 10 of differentiation (i.e. 6 days after monocyte isolation). For osteoblasts, cells were plated on day 21-28. Cells were incubated with inhibitors or vehicle at concentrations indicated in the *Compounds* section with vehicle or 10nM GIP. Media was refreshed each day. After 3 days Caspase-Glo 3/7 (Promega) and CellTiterGlo (Promega) assays were performed. Luminescence values were measured on a Pherastar FS plate reader.

### Terminal deoxynucleotidyl transferase dUTP nick-end labeling (TUNEL) staining

Cells were plated in six-well plates at 100,000 cells/well on coverslips on day 8 of differentiation. Cells were exposed to vehicle or 10nM GIP for 48 hours before fixation with 4% PFA/PBS. TUNEL staining was performed using a TUNEL FITC kit (Abcam). Cells were incubated with the staining solution for one hour, then washed and fixed in Prolong Gold Antifade reagent (Invitrogen). Images were captured using a Zeiss LSM 780 confocal microscope with a Plan-Apochromat x63/1.2/water DIC objective. An argon laser (488 nm) was used to excite Alexa Fluor 488. The raw images were then examined in ImageJ. A region of interest was selected for each nucleus, then the equivalent region examined in the FITC channel and ranked as positive or negative. Each assessor was blinded to the culture conditions.

### Real-Time PCR

RNA was extracted from osteoblasts using Trizol Plus RNA Purification Kit (Invitrogen), according to manufacturer’s instructions. The concentration and quality of total RNA were measured using NanoDrop 2000 (ThermoScientific), then 500ng of total RNA used for reverse transcription using the High-Capacity cDNA Reverse Transcription kit (Applied Biosystem). Expression levels of osteoblast specific genes (*ALPL, BGALP, COL1A1, OSX, RUNX2*) and GIPR were quantified using the Fast SYBR Green Master Mix and the Viia 7 Real-time PCR device (Applied Biosystem) and normalized to *ACTB* (β-actin). Primers used for gene expression analysis are listed in Table S1.

### Bioenergetic profiling

Osteoclasts were differentiated as described and loosened after six days exposure to RANKL. Cells were seeded at a density of 40,000 cells per well in a Seahorse 96-well plate (Seahorse XFe96, Agilent Technologies). Cells were incubated with 10nM GIP or vehicle for 3 days. Media was refreshed each day. Eight wells were seeded per donor and treatment. After 3 days, the cells were washed with seahorse assay media. Media was freshly prepared with XF DMEM Medium (Agilent Technologies) (pH 7.4) supplemented with 10 mmol/L glucose, 2 mmol/L sodium pyruvate, and 2 mmol/L glutamine (all Sigma) as previously described (*72*). Oxygen Consumption Rate (OCR) and Extracellular Acidification Rate (ECAR) were measured simultaneously in three cycles of mixing (3 min) and measuring (3 min) sequentially for each section: basal, injection 1, and injection 2. Two different groups of sequential injections were performed: (A) Injection 1: 1.5 μmol/L Oligomycin (inhibitor of complex V); Injection 2: 0.5 μmol/LM Rotenone/0.5 μM Antimycin (inhibitor of complex I and complex III, respectively); (B) Injection 1: 2.5 μmol/L FCCP (uncoupler); Injection 2: 0.5 μmol/L Rotenone/0.5 μM Antimycin. For the acute treatment with GIP, the following sequential sections were measured: basal (3 cycles), GIP injection (5 cycles), injection 1, and injection 2.

### Statistical analyses

The number of experimental replicates denoted by n is indicated in each figure legend. Data was plotted and statistical analyses performed in Graphpad Prism 7. Normality tests (Shapiro-Wilk or D’Agostino-Pearson) were performed on all datasets to determine whether parametric or non-parametric statistical tests were appropriate. A p-value of <0.05 was considered statistically significant. Statistical analyses were performed as described in figure legends but comprised either one-way ANOVA or Kruskal-Wallis with correction for multiple comparison tests, Mann-Whitney test, unpaired or nested t-tests.

## Acknowledgements

This work was funded by: Grants from the Novo Nordisk Foundation (NNF18OC0052699 (M.S.H.), NNF18OC0055047 (M.F.)), Region of Southern Denmark (ref: 18/17553 (M.S.H)) and Odense University Hospital (ref: A3147 (M.F.)), and an Academy of Medical Sciences Springboard Award supported by the British Heart Foundation, Diabetes UK, the Global Challenges Research Fund, the Government Department of Business, Energy and Industrial Strategy and the Wellcome Trust (Ref: SBF004|1034, C.M.G).

## Author contributions

M.S.H., K.S., T.L.A., M.K., C.M.G., M.F. designed experiments; C.M., J.B.O., S.O., M.M.R., B.H., R.S.H. provided materials; M.S.H., L.L.C., N.W.H., P.F.G., R.W., C.M., C.M.G. performed experiments and analyzed data. A.R. performed RNA-seq analysis. M.S.H., C.M.G., M.F. wrote the manuscript. All authors reviewed and edited the manuscript.

## References

1. J. S. Kenkre, J. Bassett, The bone remodelling cycle. Ann Clin Biochem 55, 308–327 (2018).

2. C. Holroyd, C. Cooper, E. Dennison, Epidemiology of osteoporosis. Best Pract Res Clin Endocrinol Metab 22, 671–685 (2008).

3. M. S. Hansen, M. Frost, Alliances of the gut and bone axis. Semin Cell Dev Biol, (2021).

4. D. Nasteska, N. Harada, K. Suzuki, S. Yamane, A. Hamasaki, E. Joo, K. Iwasaki, K. Shibue, T. Harada, N. Inagaki, Chronic reduction of GIP secretion alleviates obesity and insulin resistance under high-fat diet conditions. Diabetes 63, 2332–2343 (2014).

5. D. Xie, Q. Zhong, K. H. Ding, H. Cheng, S. Williams, D. Correa, W. B. Bollag, R. J. Bollag, K. Insogna, N. Troiano, C. Coady, M. Hamrick, C. M. Isales, Glucose-dependent insulinotropic peptide-overexpressing transgenic mice have increased bone mass. Bone 40, 1352–1360 (2007).

6. T. M. Kieffer, CH.; Pederson, RA., Degradation of glucose-dependent insulinotropic polypeptide and truncated glucagon-like peptide 1 in vitro and in vivo by dipeptidyl peptidase IV. Endocrinology 136, 3585–3596 (1995).

7. R. J. Bollag, Q. Zhong, P. Phillips, L. Min, L. Zhong, R. Cameron, A. L. Mulloy, H. Rasmussen, F. Qin, K. H. Ding, C. M. Isales, Osteoblast-Derived Cells Express Functional Glucose-Dependent Insulinotropic Peptide Receptors1. Endocrinology 141, 1228–1235 (2000).

8. A. Mieczkowska, B. Bouvard, D. Chappard, G. Mabilleau, Glucose-dependent insulinotropic polypeptide (GIP) directly affects collagen fibril diameter and collagen cross-linking in osteoblast cultures. Bone 74, 29–36 (2015).

9. K. Skov-Jeppesen, N. Hepp, J. Oeke, M. S. Hansen, A. Jafari, M. S. Svane, N. Balenga, J. A. Olson, Jr., M. Frost, M. Kassem, S. Madsbad, J. E. Beck Jensen, J. J. Holst, M. M. Rosenkilde, B. Hartmann, The antiresorptive effect of GIP, but not GLP-2, is preserved in patients with hypoparathyroidism-a randomized crossover study. J Bone Miner Res, (2021).

10. Q. Zhong, T. Itokawa, S. Sridhar, K. H. Ding, D. Xie, B. Kang, W. B. Bollag, R. J. Bollag, M. Hamrick, K. Insogna, C. M. Isales, Effects of glucose-dependent insulinotropic peptide on osteoclast function. Am J Physiol Endocrinol Metab 292, E543–548 (2007).

11. G. Mabilleau, R. Perrot, A. Mieczkowska, S. Boni, P. R. Flatt, N. Irwin, D. Chappard, Glucose-dependent insulinotropic polypeptide (GIP) dose-dependently reduces osteoclast differentiation and resorption. Bone 91, 102–112 (2016).

12. D. Xie, H. Cheng, M. Hamrick, Q. Zhong, K. H. Ding, D. Correa, S. Williams, A. Mulloy, W. Bollag, R. J. Bollag, R. R. Runner, J. C. McPherson, K. Insogna, C. M. Isales, Glucose-dependent insulinotropic polypeptide receptor knockout mice have altered bone turnover. Bone 37, 759–769 (2005).

13. K. Tsukiyama, Y. Yamada, C. Yamada, N. Harada, Y. Kawasaki, M. Ogura, K. Bessho, M. Li, N. Amizuka, M. Sato, N. Udagawa, N. Takahashi, K. Tanaka, Y. Oiso, Y. Seino, Gastric inhibitory polypeptide as an endogenous factor promoting new bone formation after food ingestion. Mol Endocrinol 20, 1644–1651 (2006).

14. A. Mieczkowska, N. Irwin, P. R. Flatt, D. Chappard, G. Mabilleau, Glucose-dependent insulinotropic polypeptide (GIP) receptor deletion leads to reduced bone strength and quality. Bone 56, 337–342 (2013).

15. C. Gaudin-Audrain, N. Irwin, S. Mansur, P. R. Flatt, B. Thorens, M. Basle, D. Chappard, G. Mabilleau, Glucose-dependent insulinotropic polypeptide receptor deficiency leads to modifications of trabecular bone volume and quality in mice. Bone 53, 221–230 (2013).

16. A. Nissen, M. Christensen, F. K. Knop, T. Vilsboll, J. J. Holst, B. Hartmann, Glucose-dependent insulinotropic polypeptide inhibits bone resorption in humans. J Clin Endocrinol Metab 99, E2325–2329 (2014).

17. N. C. Bergmann, A. Lund, L. S. Gasbjerg, N. R. Jørgensen, L. Jessen, B. Hartmann, J. J. Holst, M. B. Christensen, T. Vilsbøll, F. K. Knop, Separate and Combined Effects of GIP and GLP-1 Infusions on Bone Metabolism in Overweight Men Without Diabetes. J Clin Endocrinol Metab 104, 2953–2960 (2019).

18. M. B. Christensen, A. Lund, S. Calanna, N. R. Jorgensen, J. J. Holst, T. Vilsboll, F. K. Knop, Glucose-Dependent Insulinotropic Polypeptide (GIP) Inhibits Bone Resorption Independently of Insulin and Glycemia. J Clin Endocrinol Metab 103, 288–294 (2018).

19. L. S. Gasbjerg, B. Hartmann, M. B. Christensen, A. R. Lanng, T. Vilsboll, N. R. Jorgensen, J. J. Holst, M. M. Rosenkilde, F. K. Knop, GIP’s effect on bone metabolism is reduced by the selective GIP receptor antagonist GIP(3-30)NH2. Bone 130, 115079 (2020).

20. L. S. Hansen, A. H. Sparre-Ulrich, M. Christensen, F. K. Knop, B. Hartmann, J. J. Holst, M. M. Rosenkilde, N-terminally and C-terminally truncated forms of glucose-dependent insulinotropic polypeptide are high-affinity competitive antagonists of the human GIP receptor. Br J Pharmacol 173, 826–838 (2016).

21. H. S. Kizilkaya, K. V. Sorensen, C. J. Kibsgaard, L. S. Gasbjerg, A. S. Hauser, A. H. Sparre-Ulrich, N. Grarup, M. M. Rosenkilde, Loss of Function Glucose-Dependent Insulinotropic Polypeptide Receptor Variants Are Associated With Alterations in BMI, Bone Strength and Cardiovascular Outcomes. Front Cell Dev Biol 9, 749607 (2021).

22. D. M. Merrild, D. C. Pirapaharan, C. M. Andreasen, P. Kjaersgaard-Andersen, A. M. Moller, M. Ding, J. M. Delaisse, K. Soe, Pit-and trench-forming osteoclasts: a distinction that matters. Bone Res 3, 15032 (2015).

23. K. Sato, A. Suematsu, T. Nakashima, S. Takemoto-Kimura, K. Aoki, Y. Morishita, H. Asahara, K. Ohya, A. Yamaguchi, T. Takai, T. Kodama, T. A. Chatila, H. Bito, H. Takayanagi, Regulation of osteoclast differentiation and function by the CaMK-CREB pathway. Nat Med 12, 1410–1416 (2006).

24. S. H. Yoon, J. Ryu, Y. Lee, Z. H. Lee, H. H. Kim, Adenylate cyclase and calmodulin-dependent kinase have opposite effects on osteoclastogenesis by regulating the PKA-NFATc1 pathway. J Bone Miner Res 26, 1217–1229 (2011).

25. O. Destaing, A. Sanjay, C. Itzstein, W. C. Horne, D. Toomre, P. De Camilli, R. Baron, The tyrosine kinase activity of c-Src regulates actin dynamics and organization of podosomes in osteoclasts. Mol Biol Cell 19, 394–404 (2008).

26. P. Soriano, C. Montgomery, R. Geske, A. Bradley, Targeted disruption of the c-src proto-oncogene leads to osteopetrosis in mice. Cell 64, 693–702 (1991).

27. H. Abrahamsen, T. Vang, K. Tasken, Protein kinase A intersects SRC signaling in membrane microdomains. J Biol Chem 278, 17170–17177 (2003).

28. S. J. Kim, K. Winter, C. Nian, M. Tsuneoka, Y. Koda, C. H. McIntosh, Glucose-dependent insulinotropic polypeptide (GIP) stimulation of pancreatic beta-cell survival is dependent upon phosphatidylinositol 3-kinase (PI3K)/protein kinase B (PKB) signaling, inactivation of the forkhead transcription factor Foxo1, and down-regulation of bax expression. J Biol Chem 280, 22297–22307 (2005).

29. D. K. Luttrell, L. M. Luttrell, Not so strange bedfellows: G-protein-coupled receptors and Src family kinases. Oncogene 23, 7969–7978 (2004).

30. T. Miyazaki, A. Sanjay, L. Neff, S. Tanaka, W. C. Horne, R. Baron, Src kinase activity is essential for osteoclast function. J Biol Chem 279, 17660–17666 (2004).

31. S. Kim, K. Jee, D. Kim, H. Koh, J. Chung, Cyclic AMP inhibits Akt activity by blocking the membrane localization of PDK1. J Biol Chem 276, 12864–12870 (2001).

32. I. Nakamura, N. Takahashi, T. Sasaki, S. Tanaka, N. Udagawa, H. Murakami, K. Kimura, Y. Kabuyama, T. Kurokawa, T. Suda, et al., Wortmannin, a specific inhibitor of phosphatidylinositol-3 kinase, blocks osteoclastic bone resorption. FEBS Lett 361, 79–84 (1995).

33. R. E. Dolmetsch, K. Xu, R. S. Lewis, Calcium oscillations increase the efficiency and specificity of gene expression. Nature 392, 933–936 (1998).

34. Y. M. Yang, M. S. Kim, A. Son, J. H. Hong, K. H. Kim, J. T. Seo, S. I. Lee, D. M. Shin, Alteration of RANKL-induced osteoclastogenesis in primary cultured osteoclasts from SERCA2+/-mice. J Bone Miner Res 24, 1763–1769 (2009).

35. S. Yano, H. Tokumitsu, T. R. Soderling, Calcium promotes cell survival through CaM-K kinase activation of the protein-kinase-B pathway. Nature 396, 584–587 (1998).

36. J. B. Moon, J. H. Kim, K. Kim, B. U. Youn, A. Ko, S. Y. Lee, N. Kim, Akt induces osteoclast differentiation through regulating the GSK3beta/NFATc1 signaling cascade. J Immunol 188, 163–169 (2012).

37. G. Swarnkar, K. Karuppaiah, G. Mbalaviele, T. H. Chen, Y. Abu-Amer, Osteopetrosis in TAK1-deficient mice owing to defective NF-kappaB and NOTCH signaling. Proc Natl Acad Sci U S A 112, 154–159 (2015).

38. J. Lin, D. Lee, Y. Choi, S. Y. Lee, The scaffold protein RACK1 mediates the RANKL-dependent activation of p38 MAPK in osteoclast precursors. Sci Signal 8, ra54 (2015).

39. C. Bohm, S. Hayer, A. Kilian, M. M. Zaiss, S. Finger, A. Hess, K. Engelke, G. Kollias, G. Kronke, J. Zwerina, G. Schett, J. P. David, The alpha-isoform of p38 MAPK specifically regulates arthritic bone loss. J Immunol 183, 5938–5947 (2009).

40. V. Iotsova, J. Caamano, J. Loy, Y. Yang, A. Lewin, R. Bravo, Osteopetrosis in mice lacking NF-kappaB1 and NF-kappaB2. Nat Med 3, 1285–1289 (1997).

41. A. Gingery, E. Bradley, A. Shaw, M. J. Oursler, Phosphatidylinositol 3-kinase coordinately activates the MEK/ERK and AKT/NFkappaB pathways to maintain osteoclast survival. J Cell Biochem 89, 165–179 (2003).

42. S. M. Sharma, A. Bronisz, R. Hu, K. Patel, K. C. Mansky, S. Sif, M. C. Ostrowski, MITF and PU.1 recruit p38 MAPK and NFATc1 to target genes during osteoclast differentiation. J Biol Chem 282, 15921–15929 (2007).

43. M. Matsumoto, M. Kogawa, S. Wada, H. Takayanagi, M. Tsujimoto, S. Katayama, K. Hisatake, Y. Nogi, Essential role of p38 mitogen-activated protein kinase in cathepsin K gene expression during osteoclastogenesis through association of NFATc1 and PU.1. J Biol Chem 279, 45969-45979 (2004).

44. P. J. Balwierz, M. Pachkov, P. Arnold, A. J. Gruber, M. Zavolan, E. van Nimwegen, ISMARA: automated modeling of genomic signals as a democracy of regulatory motifs. Genome Res 24, 869–884 (2014).

45. S. Lemma, M. Sboarina, P. E. Porporato, N. Zini, P. Sonveaux, G. Di Pompo, N. Baldini, S. Avnet, Energy metabolism in osteoclast formation and activity. Int J Biochem Cell Biol 79, 168–180 (2016).

46. A. Mukherjee, P. Rotwein, Selective signaling by Akt1 controls osteoblast differentiation and osteoblast-mediated osteoclast development. Mol Cell Biol 32, 490–500 (2012).

47. G. Mabilleau, B. Gobron, A. Mieczkowska, R. Perrot, D. Chappard, Efficacy of targeting bone-specific GIP receptor in ovariectomy-induced bone loss. J Endocrinol, (2018).

48. M. M. Helsted, L. S. Gasbjerg, A. R. Lanng, N. C. Bergmann, S. Stensen, B. Hartmann, M. B. Christensen, J. J. Holst, T. Vilsboll, M. M. Rosenkilde, F. K. Knop, The role of endogenous GIP and GLP-1 in postprandial bone homeostasis. Bone 140, 115553 (2020).

49. M. B. N. Gabe, K. Skov-Jeppesen, L. S. Gasbjerg, S. P. Schiellerup, C. Martinussen, S. Gadgaard, G. A. Boer, J. Oeke, L. J. Torz, S. Veedfald, M. S. Svane, K. N. Bojsen-Møller, S. Madsbad, J. J. Holst, B. Hartmann, M. M. Rosenkilde, GIP and GLP-2 together improve bone turnover in humans supporting GIPR-GLP-2R co-agonists as future osteoporosis treatment. Pharmacol Res, 106058 (2022).

50. U. H. Lerner, M. Ransjo, K. Klaushofer, H. Horandner, O. Hoffmann, E. Czerwenka, K. Koller, M. Peterlik, Comparison between the effects of forskolin and calcitonin on bone resorption and osteoclast morphology in vitro. Bone 10, 377–387 (1989).

51. M. B. N. Gabe, W. J. C. van der Velden, F. X. Smit, L. S. Gasbjerg, M. M. Rosenkilde, Molecular interactions of full-length and truncated GIP peptides with the GIP receptor -A comprehensive review. Peptides 125, 170224 (2020).

52. E. A. Killion, M. Chen, J. R. Falsey, G. Sivits, T. Hager, L. Atangan, J. Helmering, J. Lee, H. Li, B. Wu, Y. Cheng, M. M. Veniant, D. J. Lloyd, Chronic glucose-dependent insulinotropic polypeptide receptor (GIPR) agonism desensitizes adipocyte GIPR activity mimicking functional GIPR antagonism. Nat Commun 11, 4981 (2020).

53. T. Miyazaki, M. Iwasawa, T. Nakashima, S. Mori, K. Shigemoto, H. Nakamura, H. Katagiri, H. Takayanagi, S. Tanaka, Intracellular and extracellular ATP coordinately regulate the inverse correlation between osteoclast survival and bone resorption. J Biol Chem 287, 37808–37823 (2012).

54. K. Skov-Jeppesen, M. S. Svane, C. Martinussen, M. B. N. Gabe, L. S. Gasbjerg, S. Veedfald, K. N. Bojsen-Møller, S. Madsbad, J. J. Holst, M. M. Rosenkilde, B. Hartmann, GLP-2 and GIP exert separate effects on bone turnover: A randomized, placebo-controlled, crossover study in healthy young men. Bone 125, 178–185 (2019).

55. R. G. Russell, Bisphosphonates: the first 40 years. Bone 49, 2–19 (2011).

56. J. P. Frías, M. J. Davies, J. Rosenstock, F. C. Pérez Manghi, L. Fernández Landó, B. K. Bergman, B. Liu, X. Cui, K. Brown, Tirzepatide versus Semaglutide Once Weekly in Patients with Type 2 Diabetes. N Engl J Med 385, 503–515 (2021).

57. P. K. Nørregaard, M. A. Deryabina, P. Tofteng Shelton, J. U. Fog, J. R. Daugaard, P. O. Eriksson, L. F. Larsen, L. Jessen, A novel GIP analogue, ZP4165, enhances glucagon-like peptide-1-induced body weight loss and improves glycaemic control in rodents. Diabetes Obes Metab 20, 60–68 (2018).

58. B. Finan, B. Yang, N. Ottaway, D. L. Smiley, T. Ma, C. Clemmensen, J. Chabenne, L. Zhang, K. M. Habegger, K. Fischer, J. E. Campbell, D. Sandoval, R. J. Seeley, K. Bleicher, S. Uhles, W. Riboulet, J. Funk, C. Hertel, S. Belli, E. Sebokova, K. Conde-Knape, A. Konkar, D. J. Drucker, V. Gelfanov, P. T. Pfluger, T. D. Muller, D. Perez-Tilve, R. D. DiMarchi, M. H. Tschop, A rationally designed monomeric peptide triagonist corrects obesity and diabetes in rodents. Nat Med 21, 27–36 (2015).

59. L. C. Hofbauer, B. Busse, R. Eastell, S. Ferrari, M. Frost, R. Muller, A. M. Burden, F. Rivadeneira, N. Napoli, M. Rauner, Bone fragility in diabetes: novel concepts and clinical implications. Lancet Diabetes Endocrinol 10, 207–220 (2022).

60. D. C. Pirapaharan, J. B. Olesen, T. L. Andersen, S. B. Christensen, P. Kjaersgaard-Andersen, J. M. Delaisse, K. Soe, Catabolic activity of osteoblast lineage cells contributes to osteoclastic bone resorption in vitro. J Cell Sci 132, (2019).

61. N. E. Lassen, T. L. Andersen, G. G. Pløen, K. Søe, E. M. Hauge, S. Harving, G. E. T. Eschen, J. M. Delaisse, Coupling of Bone Resorption and Formation in Real Time: New Knowledge Gained From Human Haversian BMUs. J Bone Miner Res 32, 1395–1405 (2017).

62. M. E. Abdelgawad, J. M. Delaisse, M. Hinge, P. R. Jensen, R. W. Alnaimi, L. Rolighed, L. H. Engelholm, N. Marcussen, T. L. Andersen, Early reversal cells in adult human bone remodeling: osteoblastic nature, catabolic functions and interactions with osteoclasts. Histochem Cell Biol 145, 603–615 (2016).

63. A. Dobin, C. A. Davis, F. Schlesinger, J. Drenkow, C. Zaleski, S. Jha, P. Batut, M. Chaisson, T. R. Gingeras, STAR: ultrafast universal RNA-seq aligner. Bioinformatics 29, 15–21 (2012).

64. S. Heinz, C. Benner, N. Spann, E. Bertolino, Y. C. Lin, P. Laslo, J. X. Cheng, C. Murre, H. Singh, C. K. Glass, Simple combinations of lineage-determining transcription factors prime cis-regulatory elements required for macrophage and B cell identities. Mol Cell 38, 576–589 (2010).

65. M. I. Love, W. Huber, S. Anders, Moderated estimation of fold change and dispersion for RNA-seq data with DESeq2. Genome Biol 15, 550 (2014).

66. M. D. Young, M. J. Wakefield, G. K. Smyth, A. Oshlack, Gene ontology analysis for RNA-seq: accounting for selection bias. Genome Biol 11, R14 (2010).

67. K. Soe, J. M. Delaisse, Glucocorticoids maintain human osteoclasts in the active mode of their resorption cycle. J Bone Miner Res 25, 2184–2192 (2010).

68. A. S. Hobolt-Pedersen, J. M. Delaisse, K. Soe, Osteoclast fusion is based on heterogeneity between fusion partners. Calcif Tissue Int 95, 73–82 (2014).

69. K. Søe, J. M. Delaissé, Time-lapse reveals that osteoclasts can move across the bone surface while resorbing. J Cell Sci 130, 2026–2035 (2017).

70. K. Viloria, D. Nasteska, L. J. B. Briant, S. Heising, D. P. Larner, N. H. F. Fine, F. B. Ashford, G. da Silva Xavier, M. J. Ramos, A. Hasib, F. Cuozzo, J. E. Manning Fox, P. E. MacDonald, I. Akerman, G. G. Lavery, C. Flaxman, N. G. Morgan, S. J. Richardson, M. Hewison, D. J. Hodson, Vitamin-D-Binding Protein Contributes to the Maintenance of alpha Cell Function and Glucagon Secretion. Cell Rep 31, 107761 (2020).

71. C. M. Gorvin, A. Rogers, B. Hastoy, A. I. Tarasov, M. Frost, S. Sposini, A. Inoue, M. P. Whyte, P. Rorsman, A. C. Hanyaloglu, G. E. Breitwieser, R. V. Thakker, AP2sigma Mutations Impair Calcium-Sensing Receptor Trafficking and Signaling, and Show an Endosomal Pathway to Spatially Direct G-Protein Selectivity. Cell Rep 22, 1054–1066 (2018).

72. P. Fernandez-Guerra, A. C. Gonzalez-Ebsen, S. E. Boonen, J. Courraud, N. Gregersen, J. Mehlsen, J. Palmfeldt, R. K. J. Olsen, L. S. Brinth, Bioenergetic and Proteomic Profiling of Immune Cells in Myalgic Encephalomyelitis/Chronic Fatigue Syndrome Patients: An Exploratory Study. Biomolecules 11, (2021).

