## Supplemental Appendix for "GIP receptor reduces osteoclast activity and improves osteoblast survival by activating multiple signaling pathways"

**Supplementary Appendix**

**Figure S1	GIP reduces bone resorption and increases cAMP in osteoclast monocultures and osteoclast-osteoblast co-cultures**

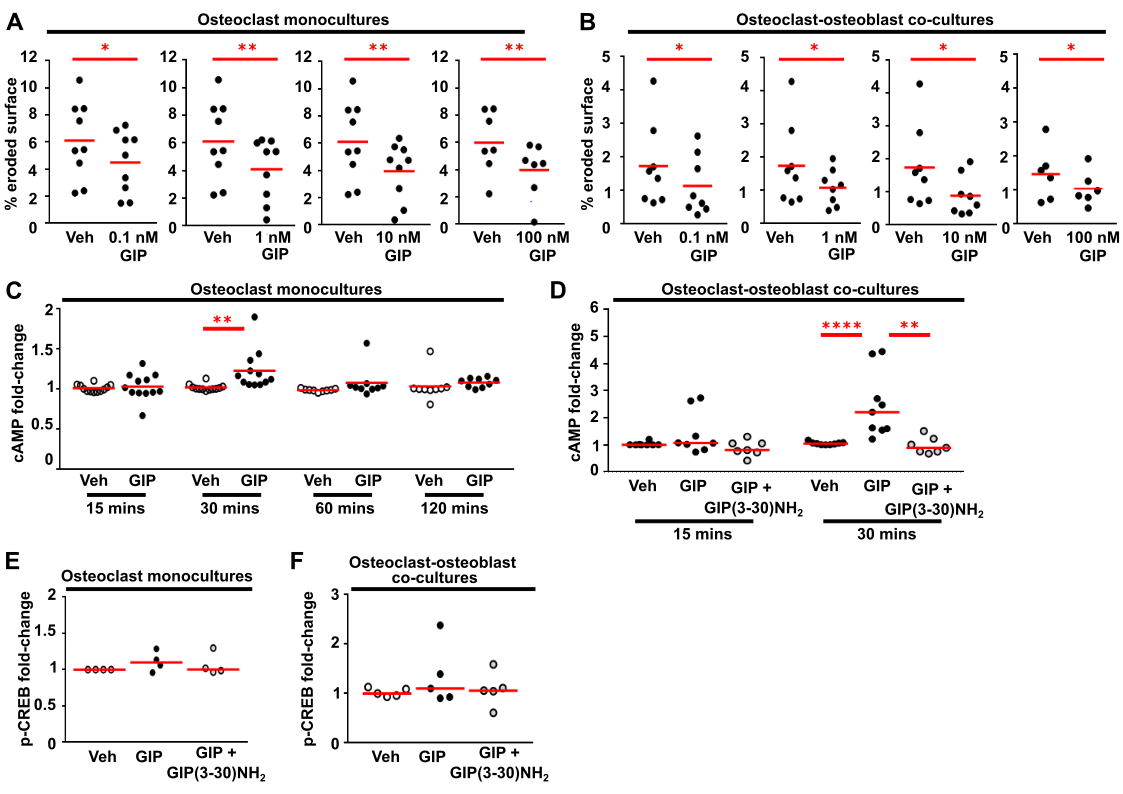

(**A-B**) Quantification of percentage eroded surface of bone slices by (A) osteoclast monocultures and (B) osteoclast-osteoblast co-cultures at four concentrations of GIP, measured using toluidine blue staining. N=6-8 donors for osteoclast monocultures and N=6-8 donors for osteoclast-osteoblast co-cultures. (**C-D**) Quantification of cAMP generated in (C) osteoclasts and (D) osteoclast-osteoblast co-cultures measured by LANCE assays. N=9-12 for osteoclasts and N=7-9 for osteoclast-osteoblast co-cultures. (**E-F**) Quantification of phosphorylated CREB (p-CREB) in (E) osteoclasts and (F) osteoclast-osteoblast co-cultures following 30 minutes exposure to vehicle, 10nM GIP, or GIP and the GIPR antagonist, GIP(3-30)NH_2_, measured by AlphaLISA assay. p-CREB was normalized to GAPDH in AlphaLISA assays. N=4 donors for monocultures and N=5 for osteoclast-osteoblast co-cultures. Panels S1A and S1B were analyzed using a paired t-test; panels S1C, S1D, S1E, S1F were analyzed using Kruskal-Wallis one-way ANOVA with Dunn’s multiple comparisons test.

**Figure S2	GIP-mediated reductions in p-Src do not involve Ca^2+^_I_, PKC or PI3K-Akt signaling pathways**

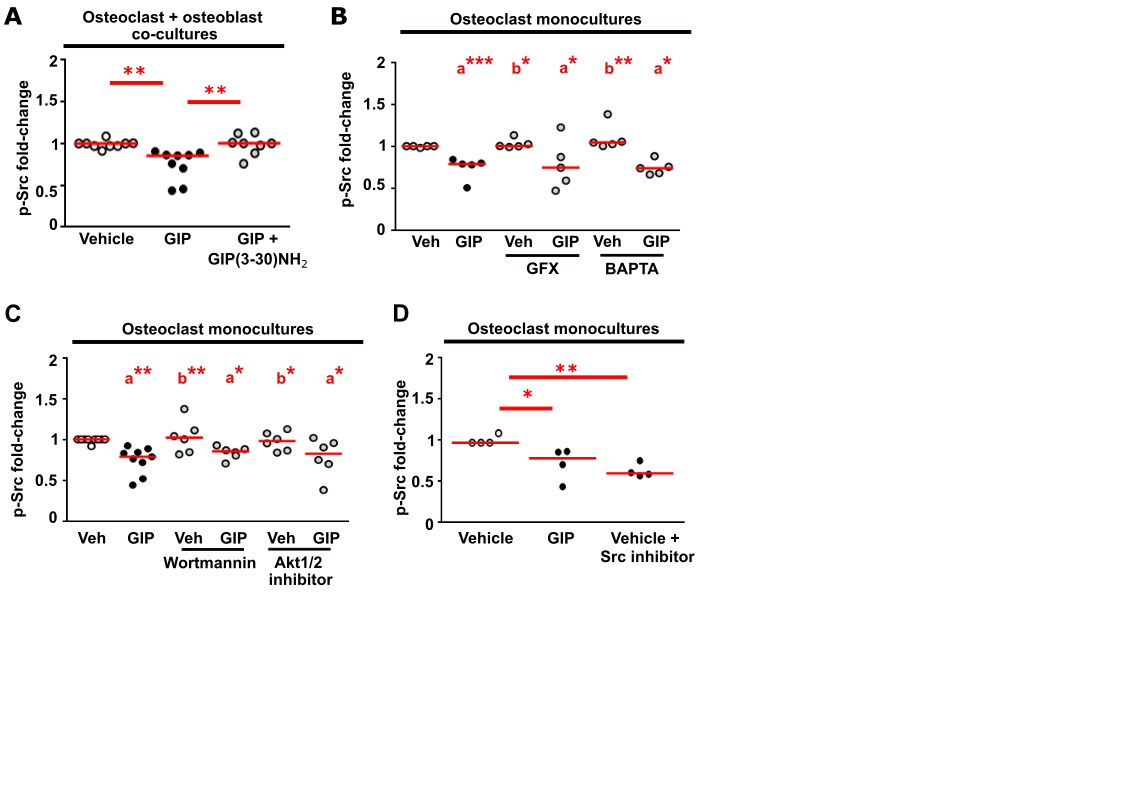

(**A**) Quantification of phosphorylated c-Src (p-Src) in osteoclast-osteoblast co-cultures following 30 minutes exposure to vehicle, GIP, or GIP and the GIPR antagonist, GIP(3-30)NH_2_, measured by AlphaLISA assay. N=8-9 donors. (**B**-**D**) Effect of pre-treatment with (B) the PKC inhibitor, GF109203X (GFX), or the calcium chelator BAPTA, (C) the PI3K inhibitor wortmannin or an Akt1/2 inhibitor, (D) a Src inhibitor, on GIP-mediated p-Src responses in osteoclast monocultures. N=5 for B, N=6 for C, N=4 donors for D. p-Src was normalized to GAPDH in all panels. Statistical comparisons to vehicle-treated cells are labelled as a, and to GIP-treated cells as b. **p<0.01, *p<0.05. Statistical analyses were performed using: Kruskal-Wallis one-way ANOVA with Dunn’s multiple comparisons test for all panels.

**Figure S3	GIP reduces p-Akt signaling in primary human osteoclast-osteoblast co-cultures**

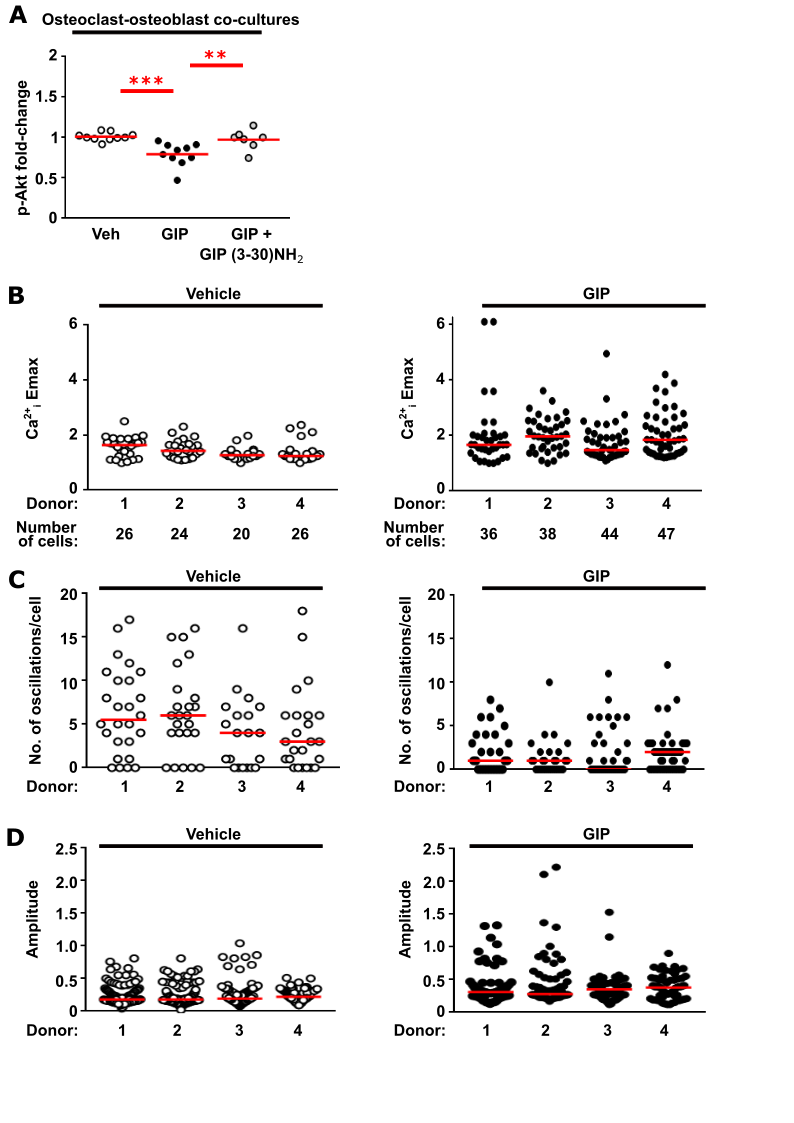

(**A**) Quantification of phosphorylated Akt1/2/3 (p-Akt) generated in osteoclast-osteoblast co-cultures by 30 minutes exposure to vehicle (veh), GIP, or GIP with the GIPR antagonist GIP(3-30)NH_2_, measured by AlphaLISA assay. N=7-10 donors. p-Akt was normalized to GAPDH. (**B**) Ca^2+^_i_ Emax for individual cells measured for each donor. There was no significant difference between the datasets for the four donors, and therefore data was combined in Figure 3G-H. (**C**) Number of oscillations in each of the cells shown in B for each donor. There was no significant difference between the datasets for the four donors, and therefore data was combined in Figure 3J-K. (**D**) Amplitude of all oscillations for each donor. There was no significant difference between the datasets for the four donors, and therefore data was combined in Figure 3L-M. ***p<0.001, **p<0.01. Panel S3A analyzed by one-way ANOVA with Tukey’s multiple comparison test. Panels S3B-S3D were analyzed using Kruskal-Wallis one-way ANOVA with Dunn’s multiple comparisons test.

**Figure S4	GIP reduces NFATc1 nuclear translocation in primary human osteoclasts**

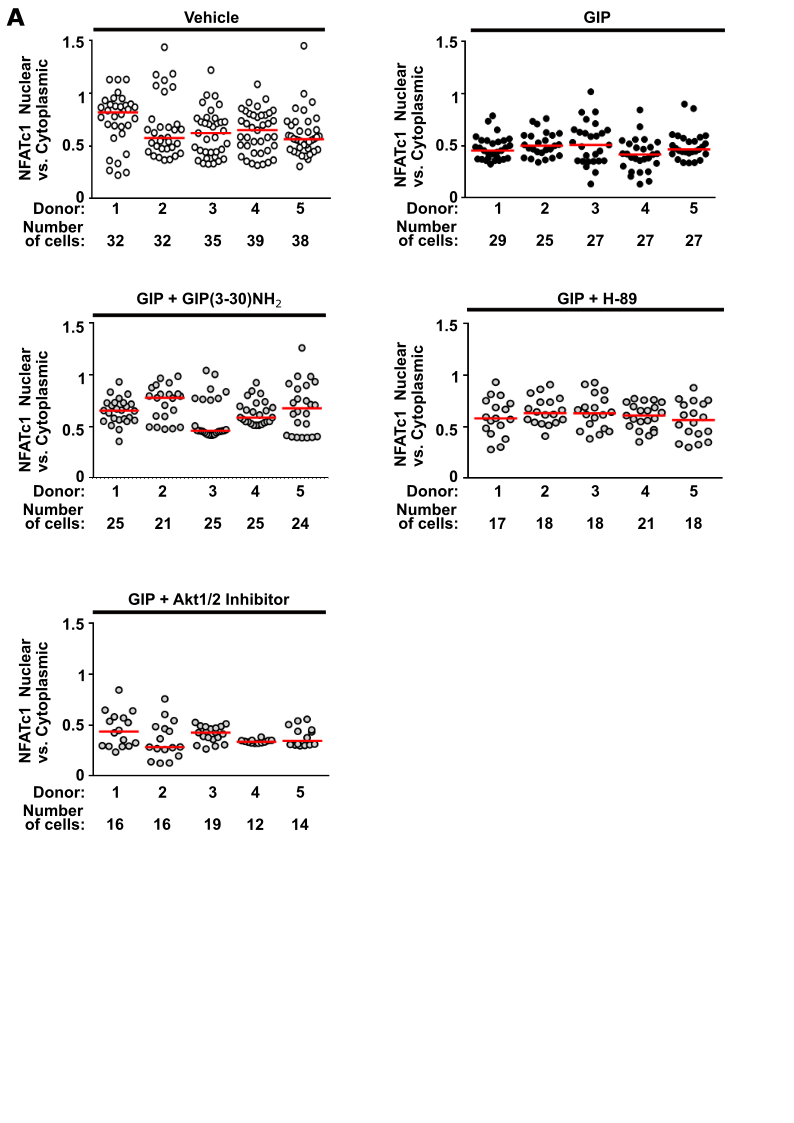

NFATc1 nuclear and cytoplasmic ratios in individual cells measured for each donor. There was no significant difference between the datasets for the four donors, therefore data was combined in Figure 4. Statistical analyses were performed using Kruskal-Wallis one-way ANOVA with Dunn’s multiple comparisons test.

**Figure S5	GIP reduces p38 and NFκB signaling in primary human osteoclasts and osteoclast-osteoblast co-cultures**

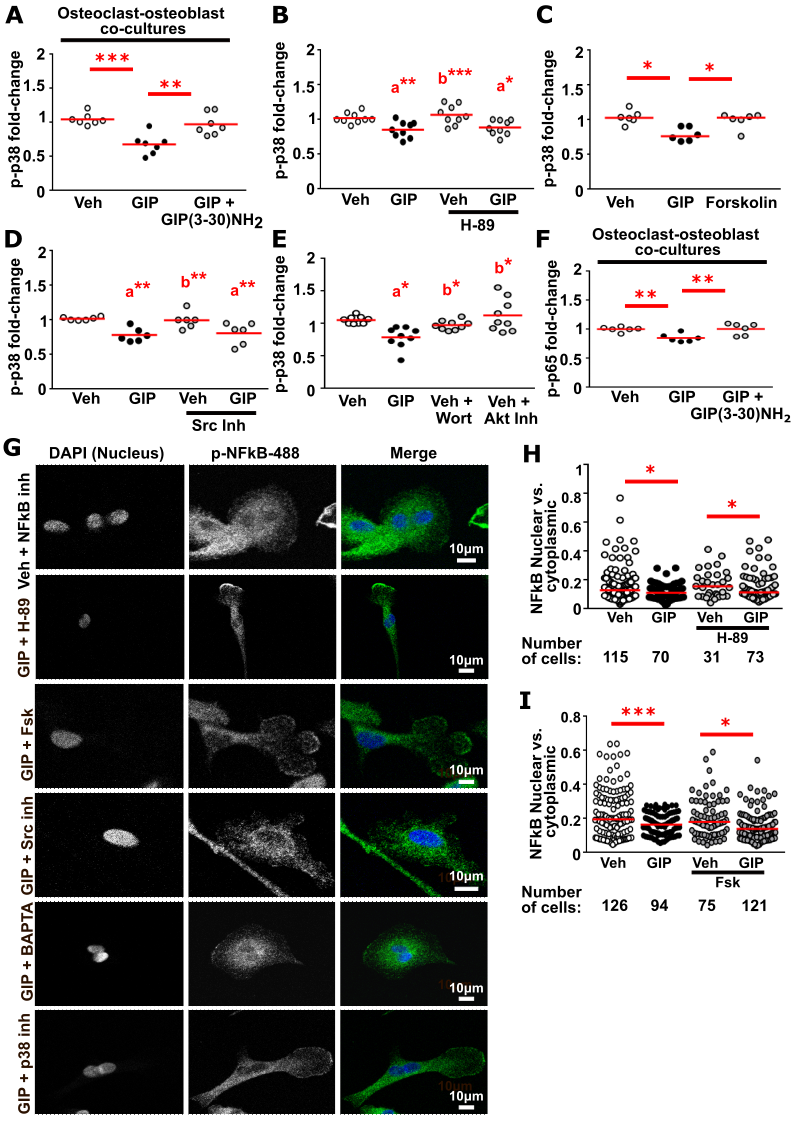

(**A**) Quantification of phosphorylated p38 (p-p38) generated in osteoclast-osteoblast co-cultures by 30 minutes exposure to vehicle (veh), GIP, or GIP with the GIPR antagonist GIP(3-30)NH_2_ measured by AlphaLISA assay. N=7 donors. (**B-E**) Effect of pre-treatment with (B) the PKA inhibitor H-89, (C) the adenylate cyclase activator Forskolin (Fsk), (D) a Src inhibitor, (E) inhibitors of PI3K (wortmannin, wort) and Akt1/2, on p-p38 in osteoclast monocultures. N=9 for B, E and N=6 donors for C, D. (**F**) Quantification of phosphorylated p65 (p-p65) generated in osteoclast-osteoblast co-cultures by 30 minutes exposure to vehicle (veh), GIP, or GIP with GIP(3-30)NH_2_, measured by AlphaLISA assay. N=6 donors. Concentrations of p-p38 and p-p65 were normalized to GAPDH. Comparisons to vehicle-treated cells are labelled as a, and to GIP-treated cells as b. ***p<0.001, **p<0.01, *p<0.05. (**G**) Representative images of p-NFκB in osteoclasts following 60 minutes exposure to vehicle or GIP +/- inhibitors of signaling. DAPI was used to label nuclei, and AlexaFluor488 secondary antibody to fluorescently label p-NFκB. (**H-I**) Quantification of the ratio of p-NFκB in nuclear and cytoplasmic fractions of cells exposed to vehicle and GIP, and pre-treated with (H) the PKA inhibitor H-89 and (I) the adenylate cyclase activator forskolin (Fsk). Statistical analyses were performed by: one-way ANOVA with Dunnett’s test for panels S5A, S5B, S5D, S5E, S5F; and Kruskal-Wallis one-way ANOVA with Dunn’s test for panels S5G and S5H.

**Figure S6	GIP reduces NFκB signaling in human osteoclasts**

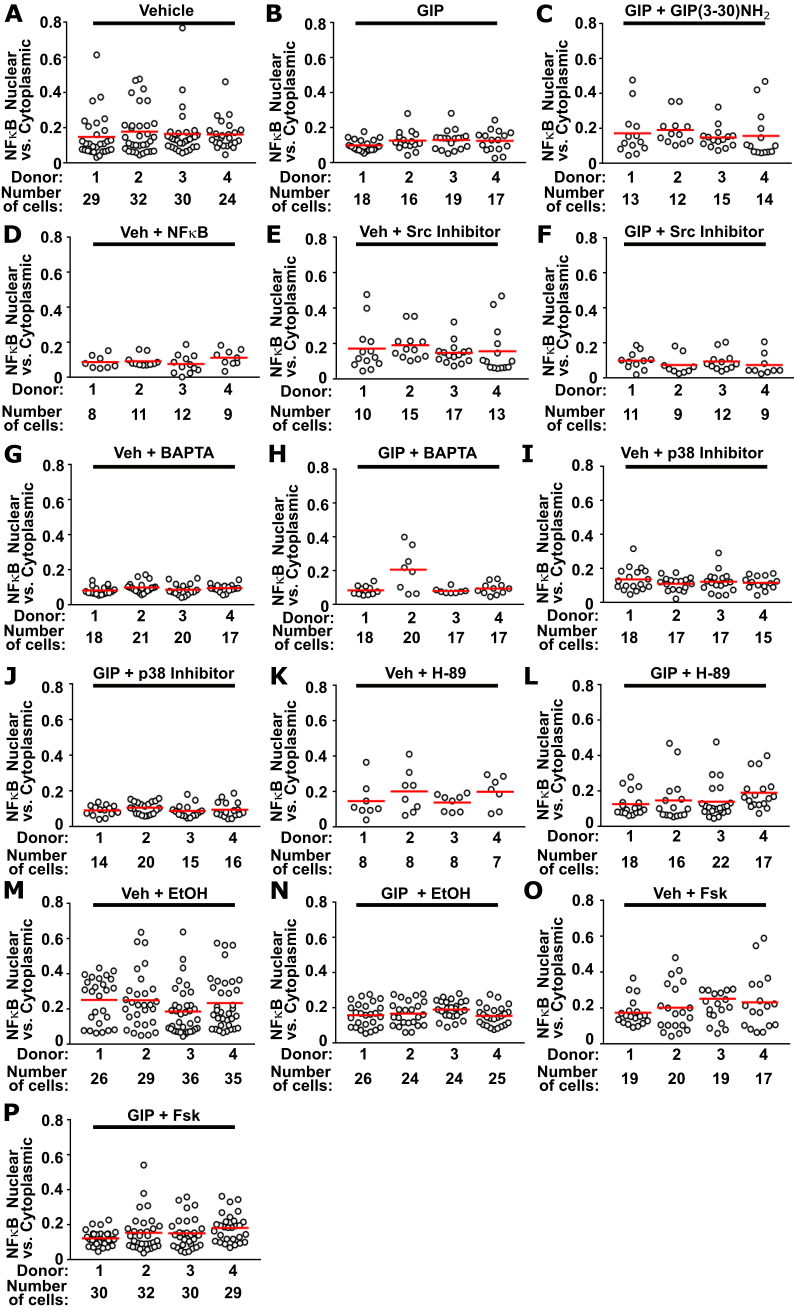

(**A**-**P**) p-NFκB nuclear and cytoplasmic ratios in osteoclasts from N=4 donors. Data for all cells measured for individual donors is shown.

**Figure S7	GIP increases cAMP in human osteoclasts and osteoblasts**

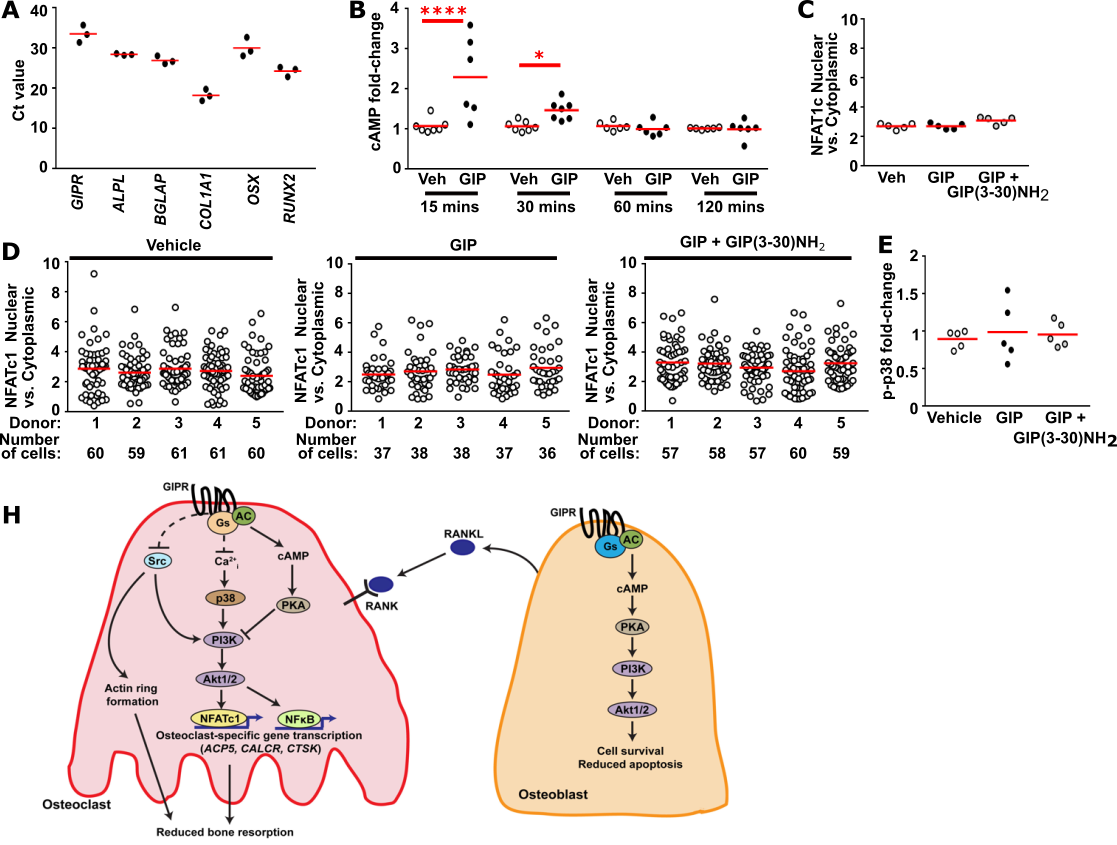

(**A**) Cycle threshold (Ct) values for *GIPR* and genes with important osteoblast functions assessed by qPCR. N=3 donors. *ALPL*, alkaline phosphatase; *BGLAP*, bone gamma-carboxyglutamate protein (osteocalcin gene); *COLA1A*, collagen type 1, alpha 1 chain; *OSX*, osterix; *RUNX2*, Runt-related transcription factor 2. (**B**) Quantification of cAMP in osteoblasts exposed to vehicle, GIP or GIP+GIP(3-30)NH_2_ for 0, 15, 30, 60 and 120 minutes measured by LANCE assays. Data is normalized to cAMP at time 0. N=6-7 donors. (**C-D**) NFATc1 nuclear and cytoplasmic ratios in osteoclasts from N=5 donors. Panel C shows average for each of the four donors. Panel D-F shows data for individual donors. (**E**) Quantification of phosphorylated p-38 in osteoblasts following 30 minutes exposure to vehicle or 10nM GIP measured by AlphaLISA assay. Data was normalized to GAPDH in AlphaLISA assays. N=5 donors. (**F**) GIPR activates canonical Gs-mediated cAMP-PKA signaling in osteoclasts to reduce NFATc1 nuclear translocation and osteoclast-specific gene transcription. GIPR also inhibits Src phosphorylation to reduce actin ring formation, and impairs intracellular calcium signaling (Ca^2+^_I_) to reduce p38-PI3K-Akt signaling and reduce NFATc1 and NFκB nuclear translocation and gene transcription. Combined actions of GIP on these signaling pathways reduces bone resorption. GIP activation of GIPR on osteoblasts activates cAMP-PKA and PI3K-Akt signaling to reduce apoptosis and improve cell survival. GIP actions on osteoclasts rely on intact RANKL-RANK signaling between osteoblasts and osteoclasts. Abbreviations: AC, adenylate cyclase; ACP5, acid phosphatase type 5 (TRAP) gene; CALCR, calcitonin receptor gene; CTSK, cathepsin K gene; Gs, stimulatory G protein; NFATc1, nuclear translocation of nuclear factor of activated T cells 1; NFκB, nuclear factor-κB; PI3K, phosphatidylinositol 3-kinase; PKA, protein kinase A; RANKL, receptor activator of NFκB ligand. Panels S7B and S7D were analyzed by Kruskal-Wallis one-way ANOVA with Dunn’s test. Panels S7C and S7E were analyzed by one-way ANOVA with Tukey’s multiple comparison test.

**Table S1	List of SYBR green primers**

| Gene name | Sequence |
| --- | --- |
| *ALPL* | Forward: ACGTGGCTAAGAATGTCATC  Reverse: CTGGTAGGCGATGTCCTTA |
| *BGLAP* | Forward: CGCTACCTGTATCAATGGCTGG  Reverse: CTCCTGAAAGCCGATGTGGTCA |
| *COL1A1* | Forward: AGGGCTCCAACGAGATCGAGATCCG  Reverse: TACAGGAAGCAGACAGGGCCAACGTCG |
| *GIPR* | Forward: CTGCCTGCCGCACGGCCCAGAT  Reverse: GCGAGCCAGCCTCAGCCGGTAA |
| *OSX* | Forward: AACCCCCAGCTGCCCACCTACC  Reverse: ACTGCCCCCATATCCACCACTACCC |
| *RUNX2* | Forward: TGGTTACTGTCATGGCGGGTA  Reverse: TCTCAGATCGTTGAACCTTGCTA |

**Table S2	Effects of 30 minutes exposure to GIP on the bioenergetics profile of human osteoclasts**

|  |  | **GIP (mean ± SEM)** | | | | | | | | |
| --- | --- | --- | --- | --- | --- | --- | --- | --- | --- | --- |
| **Mitochondrial related parameters** | | **0 nM** |  | **0.1 nM** |  | **1 nM** |  | **10 nM** |  | **100 nM** |
|  | **Basal respiration**  **(pmol oxygen/min)** | 60.50 ± 5.25 |  | 68.94 ± 4.80 |  | 72.27 ± 5.01 |  | 66.43 ± 3.88 |  | 63.80 ± 3.67 |
|  | **Induced basal respiration (pmol oxygen/min)** | 61.33 ± 6.58 |  | 70.45 ± 5.85 |  | 72.72 ± 6.29 |  | 66.70 ± 5.19 |  | 62.86 ± 4.81 |
|  | **Proton leak**  **(pmol oxygen/min)** | 9.52 ± 1.50 |  | 10.93 ± 1.57 |  | 11.17 ± 1.85 |  | 10.98 ± 1.61 |  | 10.21 ± 1.40 |
|  | **ATP-linked respiration**  **(pmol oxygen/min)** | 51.81 ± 5.08 |  | 59.51 ± 4.29 |  | 61.55 ± 4.47 |  | 55.73 ± 3.62 |  | 52.66 ± 3.43 |
|  | **Non-mitochondrial oxygen consumption**  **(pmol oxygen/min)** | 20.62 ± 2.58 |  | 22.48 ± 2.53 |  | 23.80 ± 2.61 |  | 22.68 ± 2.15 |  | 22.04 ± 2.01 |
|  | **Coupling efficiency**  **(%)** | 85.37 ± 1.07 |  | 86.29 ± 0.41 |  | 85.10 ± 0.31 |  | 83.80 ± 0.57 |  | 82.42 ± 0.68 |
|  |  | **GIP (mean ± SEM)** | | | | | | | | |
| **ATP rate related parameters** | | **0 nM** |  | **0.1 nM** |  | **1 nM** |  | **10 nM** |  | **100 nM** |
|  | **Mitochondrial ATP production rate (pmol/min)** | 287.33 ± 28.03 |  | 328.75 ± 23.60 |  | 339.95 ± 25.14 |  | 308.85 ± 20.86 |  | 291.45 ± 18.94 |
|  | **Glycolytic ATP production rate (pmol/min)** | 84.90 ± 26.64 |  | 106.54 ± 36.88 |  | 119.49 ± 40.95 |  | 118.47 ± 43.20 |  | 110.86 ± 34.20 |
|  | **Total ATP production rate (pmol/min)** | 372.20 ± 54.24 |  | 435.30 ± 60.43 |  | 459.43 ± 65.91 |  | 427.33 ± 63.44 |  | 402.30 ± 52.93 |
|  | **ATP rate index (mitochondrial ATP/glycolytic ATP)** | 4.59 ± 1.09 |  | 4.74 ± 1.39 |  | 4.22 ± 1.24 |  | 4.16 ± 1.35 |  | 3.61 ± 0.91 |
|  | **% Glycolysis** | 20.74 ± 3.94 |  | 21.94 ± 5.19 |  | 23.49 ± 5.48 |  | 24.77 ± 6.36 |  | 25.50 ± 5.03 |
|  | **% Oxidative phosphorylation** | 79.26 ± 3.94 |  | 78.06 ± 5.19 |  | 76.51 ± 5.48 |  | 75.24 ± 6.36 |  | 74.50 ± 5.03 |

Bioenergetic profile of human osteoclasts following treatment with vehicle or four concentrations of GIP (0.1, 1, 10, 100 nM) for 30 minutes. Data shows mean±SEM (n = 4). All parameters were analyzed by paired t-test and p-values <0.05 were considered significant. Following multiple-comparisons testing, there were no significant differences between any parameters.

**Table S3	Effects of 3 day exposure to GIP on the bioenergetics profile of human osteoclasts**

|  |  | **GIP (mean ± SEM)** | | |  |  |  |  |
| --- | --- | --- | --- | --- | --- | --- | --- | --- |
| **Mitochondrial related parameters** | | **0 nM** |  | **10 nM** |  | **Fold-change** |  | **p-value** |
|  | **Basal respiration**  **(pmol oxygen/min)** | 42.58 ± 5.04 |  | 52.58 ± 5.15 |  | 1.23 |  | **0.01** |
|  | **Proton leak**  **(pmol oxygen/min)** | 12.92 ± 0.85 |  | 14.10 ± 1.08 |  | 1.09 |  | 0.24 |
|  | **ATP-linked respiration**  **(pmol oxygen/min)** | 39.75 ± 4.57 |  | 46.20 ± 5.31 |  | 1.16 |  | 0.13 |
|  | **Maximal respiration**  **(pmol oxygen/min)** | 251.32 ± 32.44 |  | 297.03 ± 24.73 |  | 1.18 |  | **0.04** |
|  | **Spare respiratory capacity (pmol oxygen/mint** | 3.87 ± 0.32 |  | 3.94 ± 0.26 |  | 1,02 |  | 0.68 |
|  | **Non-mitochondrial oxygen consumption**  **(pmol oxygen/min)** | 37.07 ± 1.66 |  | 35.33 ± 1.80 |  | 0.95 |  | 0.10 |
|  | **Coupling efficiency**  **(%)** | 93.94 ± 5.36 |  | 87.22 ± 2.81 |  | 0.93 |  | 0.27 |
|  | | **GIP (mean ± SEM)** | | |  |  |  |  |
| **ATP rate related parameters** | | **0 nM** |  | **10 nM** |  | **Fold-change** |  | **p-value** |
|  | **Mitochondrial ATP production rate**  **(pmol/min)** | 223.40 ± 26.05 |  | 255.40 ± 30.54 |  | 1.14 |  | 0.12 |
|  | **Glycolytic ATP production rate**  **(pmol/min)** | 250.10 ± 62.95 |  | 246.70 ± 40.15 |  | 0.99 |  | 0.73 |
|  | **Total ATP production rate**  **(pmol/min)** | 473.50 ± 76.17 |  | 544.15 ± 63.72 |  | 1.15 |  | 0.71 |
|  | **ATP rate index (mitochondrial ATP/glycolytic ATP)** | 1.07 ± 0.16 |  | 1.13 ± 0.10 |  | 1.06 |  | 0.45 |
|  | **% Glycolysis** | 49.08 ± 4.74 |  | 47.46 ± 2.65 |  | 0.97 |  | 0.26 |
|  | **% Oxidative phosphorylation** | 50.93 ± 4.74 |  | 52.54 ± 2.65 |  | 1.03 |  | 0.26 |

Bioenergetic profile of human osteoclasts following treatment with vehicle or 10nM GIP for 3 days. Significant p-values are shown in bold (n = 6). All parameters were analyzed by paired t-test and p-values <0.05 were considered significant. To adjust for multiple comparisons, we established a fold-change criterion higher than 20% to be considered biologically relevant.
